# Artificial Intelligence of Things-Enhanced Automated Surveillance System for Global Antimicrobial Resistance in Food Supply Chain

**DOI:** 10.64898/2026.02.10.705121

**Authors:** Jinxin Liu, Marti Z. Hua, Xinyu Yan, Luyao Ma, Shenmiao Li, Yang Wang, Tian Yang, Yihan He, Michael E. Konkel, Greta Gölz, Thomas Alter, Jinsong Feng, Qian Liu, Xiaonan Lu

## Abstract

Antimicrobial resistance (AMR) threatens food safety across the farm-to-fork continuum. Real-time surveillance is crucial to mitigate its global escalation, yet conventional antimicrobial susceptibility testing (AST) remains slow, labor-intensive, and impractical for large-scale monitoring. We developed an Artificial Intelligence of Things (AIoT)-integrated multiplex microfluidic platform enabling automated AMR surveillance of pathogens in food supply chain. Each node combines a single-board AIoT controller (Orange Pi 5B), portable incubator, colorimetric microfluidic chips, and environmental sensors, reducing costs by 98% compared with standard AST. A lightweight YOLOv5 model embedded in the controller achieved >99% accuracy in identifying bacterial growth and inhibition under antibiotic pressure, showing 96% and 95% agreement with standard results for *Salmonella* and *Campylobacter*, respectively. Data are synchronized to a cloud server for real-time aggregation and early resistance warning. This fully automated and low-cost system minimizes human error and workload, providing a scalable sample-to-answer solution for AMR surveillance in global agri-food system.

## 1. Main

The World Health Organization (WHO) has recognized antimicrobial resistance (AMR) as one of the top three global health threats ^1^. The effectiveness of antibiotics against infectious diseases has been diminishing, exacerbated by the lack of discovery and development of innovative antibiotic drugs since the 1970s ^2,3^. AMR is associated with nearly 700,000 deaths annually and is projected to cause up to 10 million deaths per year by 2050 if not addressed ^4^. There is a universal consensus on the need for continuous global surveillance of AMR to monitor disease trends, guide public health strategies, inform treatment decisions, and support research and policy development. Given the significant role of the global food supply chain in rapid spread of multidrug-resistant pathogenic bacteria ^5^, it is particularly vital for the agri-food sector to proactively predict and mitigate AMR threats.

Substantial progress has been made in tracking the spread of AMR. Over the past decades, various surveillance systems have been established to monitor AMR trends in humans, animals and food. For example, the Canadian Antimicrobial Resistance Surveillance System (CARSS) collects data on antimicrobial use and resistance in humans and animals to inform public health decisions. In the US, the National Antimicrobial Resistance Monitoring System (NARMS) monitors antibiotic use and resistance in meat and poultry products. Denmark’s DANMAP oversees antibiotic usage and resistance across food-producing animals, food, feed, and humans. The RESAPATH network in France tracks antimicrobial resistance in pathogens from food-producing animals. The WHO’s Global Antimicrobial Resistance and Use Surveillance System (GLASS) urges member countries to adopt standardized protocols for collecting and analyzing AMR data ^6^. However, significant challenges remain in consolidating data across regions, resulting in fragmented and inconsistent data sharing. Combined with the time-intensive and resource-demanding nature of traditional antimicrobial susceptibility testing (AST) that requires high-standard laboratory facilities and skilled personnel, these limitations indicate that AMR surveillance trends are typically reported only on an annual basis, restricting timely risk assessment and intervention.

Microfluidic lab-on-a-chip technology offers distinct advantages over traditional macroscale approaches, including cost efficiency, high portability, and reduced manual labor, making it particularly valuable for rapid and on-site screening of foodborne pathogens ^7^. These miniaturized devices manipulate small fluid volumes (e.g., on a microliter or nanoliter scale) within tiny channels, enabling rapid analysis, cost efficiency, and ease of operation ^8^. Microfluidic-based AST has been developed to rapidly determine bacterial susceptibility based on either morphological or physiological changes. Analysis of bacterial growth within microfluidic channels is typically performed using phase contrast microscopy, fluorescent microscopy, or visible colorimetric reactions ^9,10^. However, manual analysis of sensing data can be inaccurate and slow due to signal noise and human error. To address these challenges, machine learning algorithms can be integrated with conventional sensors to create intelligent sensors that automatically predict bacterial profiles using decision systems. For example, Coelho and colleagues applied a decision tree model to 1,632 *Staphylococcus aureus* isolates incorporating minimum inhibitory concentration (MIC) and minimum bactericidal concentration (MBC) values for four biocides, together with collection year, geographic origin, host demographics, and infection details, to predict antibiotic susceptibility ^11^. In another study, Pyayt and colleagues used a random forest classifier on time- and angle-resolved scattered-light microscopy data from single bacterial cells, extracting growth- and division-related features to rapidly assess antibiotic susceptibility ^12^. Moreover, Thrift and colleagues applied a variational autoencoder to distinguish antibiotic responses of *Escherichia coli* and *Pseudomonas aeruginosa* using surface-enhanced Raman spectroscopy. Trained on large-scale spectral datasets, the model was able to extract complex spectral features associated with bacterial responses, including subtle signals that are not readily detectable by human experts ^13^. Although these algorithms significantly improved detection accuracy and speed, numerous manual experiments are still required before data can be collected and transferred to the computer for algorithm processing. Further automation of data collection process will be essential to alleviate this bottleneck and maximize the efficiency of microfluidic-based AST systems.

The Internet of Things (IoT) is a network of sensor-embedded devices capable of communication and collaboration ^14^. It has been used in industrial automation ^15^, precision farming ^16^, and smart cities ^17,18^. These IoT-enabled devices offer real-time monitoring and response capabilities, enabling immediate actions and long-term surveillance ^19,20^. In the food industry, the automation features of IoT minimizes human error and enhances efficiency. For example, Wang and co-authors incorporated radio frequency identification (RFID) technology to monitor individual perching behavior in group-housed poultry ^21^. In another study, Cocco and others used Raspberry Pi to record temperature and humidity data on a blockchain, providing consumers transparent insights into the entire trajectory from the post-harvest raw materials to the final product ^22^. Moreover, Sourav and colleagues developed a smart system using four Arduino microcontrollers with connected sensors to monitor perishable food during storage ^23^. The successful applications of IoT in the agri-food industry demonstrate its potential for AMR detection and surveillance in the agroecosystem and food supply chain. In a previous study, Ma and others developed a prototype AMR surveillance system using IoT sensors and microfluidic “lab-on-a-chip” ^24^. It incorporated a portable incubator for bacterial cultivation, real-time camera-laptop monitoring, and machine learning-based AMR classification. While efficient and cost-effective, it faced scalability limitations. In addition, it was evaluated using a single bacterial species and could process only one bacterial target at a time without multiplexing capability. Moreover, this system relied on costly GPUs for machine learning.

In this study, we developed an automated AMR surveillance system that integrates multiplex microfluidics with Artificial Intelligence of Things (AIoT), which is a paradigm that combines artificial intelligence (AI) with IoT connectivity to enable autonomous data acquisition, real-time analysis, and remote control. We focused on *Campylobacter* and *Salmonella*, two major bacterial pathogens prevalent in the agri-food system and recognized as leading causes of foodborne illnesses in humans. Designed for high-priority applications in the food supply chain, our system enables simultaneous detection of both pathogens and assessment of their resistance to antibiotics critical for clinical therapy and AMR surveillance. *Salmonella* was tested against ampicillin, ciprofloxacin, erythromycin, and tetracycline, representing distinct antimicrobial classes used in global monitoring programs. *Campylobacter* was tested against ciprofloxacin, erythromycin, and tetracycline, which are essential for clinical management and frequently reported in public health surveillance due to their high resistance rates. The microfluidic chip cultivates target pathogens in chromogenic agar to allow visual differentiation. An IoT-connected camera continuously captures images of the microfluidic chips, which are processed in real time by an AI model running on the AIoT device (Orange Pi 5B) to analyze color changes and determine AMR phenotypes without manual input. Results are then transmitted to a cloud server for further processing. Leveraging the built-in computational power (6 TOPS) of the AIoT device, this system avoids reliance on costly GPUs, keeping hardware costs under USD $150 excluding the portable incubator of ∼USD $20 that is scalable to different detection volumes. This durable AIoT hardware supports long-term and repeated monitoring. By combining affordability, automation, scalability, and reusability, it offers a practical solution for globally integrated AMR surveillance in the food supply chain particularly in resource-limited settings.

## 2. Results

### 2.1. Architecture of the global AMR surveillance system

Our next-generation AMR surveillance system is designed to improve detection efficiency and accuracy through an AIoT-integrated workflow that links sensor-based data acquisition with real-time and on-chip bacterial testing, thereby minimizing errors associated with manual operations. The proposed AMR surveillance system adopts a three-layer architecture, including perception layer, transport layer, and application layer (**Fig. 1**). The foundation is the perception layer that hosts numerous intelligent edge nodes distributed across various detection terminals, such as fields, retail stores, and food production facilities. An edge node is a device or endpoint located near or at the boundary of the network that collects, processes, and communicates data, serving as the primary interaction point with the physical world ^25^. Each edge node rapidly completes most functional tasks without uploading raw data to the cloud. In this AIoT system, an AI-enhanced embedded device called Orange Pi 5B serves as the intelligent edge node. It monitors and controls temperature, humidity, oxygen, and CO_2_ levels within the portable incubator. Additionally, a high-resolution camera connected to Orange Pi observes the colorimetric patterns of the microfluidic chip via the glass lid of the incubator. A lightweight detection model compressed from YOLO v5 detects individual wells in each microfluidic chip. Each intelligent edge node connects to the internet to form an IoT network known as the transport layer, which manages transmission strategies, resolves congestion issues, and ensures data security and integrity. Finally, the application layer that is typically hosted on a cloud computing platform interacts directly with users. After aggregating and analyzing data from the edge nodes, it offers advanced features such as AMR identification, extraction of temporal and spatial trends, early warning generation, and algorithm updates that are deployed back to the edge nodes.

**Fig. 1.**
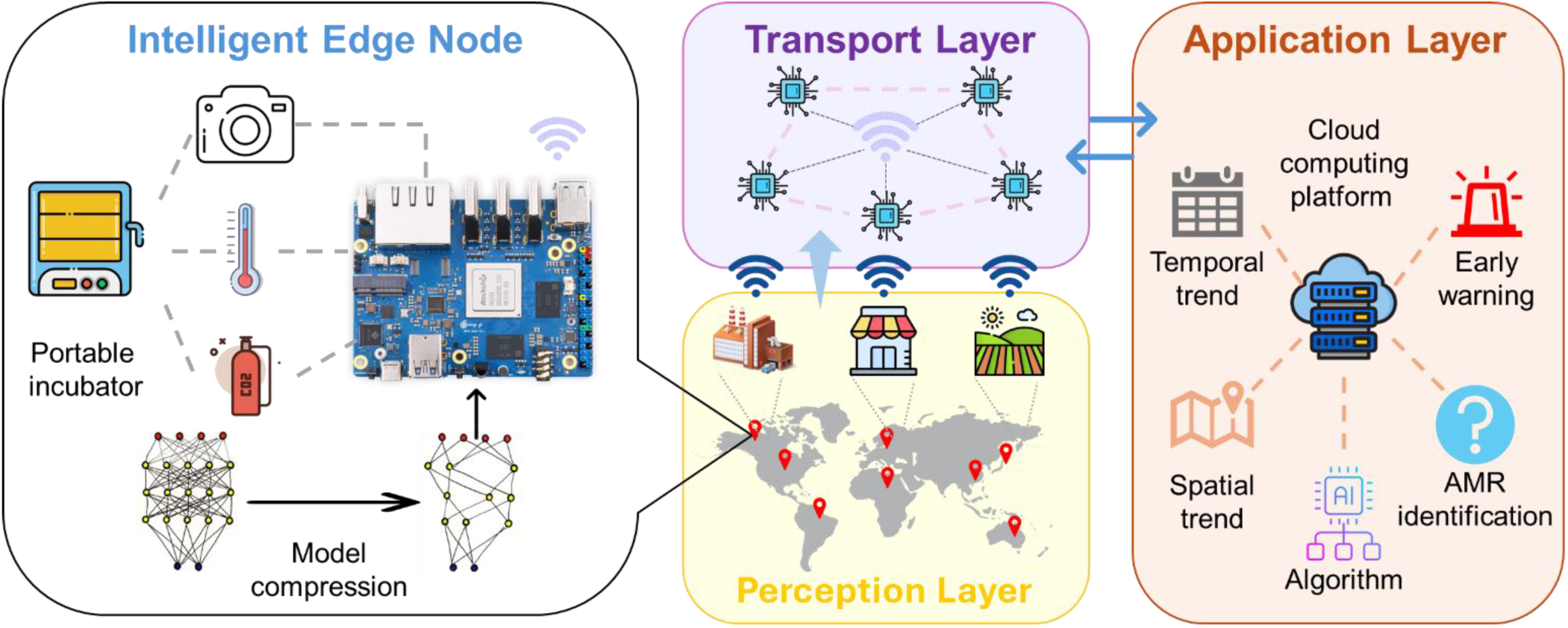
Schematic overview of a global AMR surveillance system integrating embedded AI and microfluidic technology. The system is structured into three primary layers: perception, transport, and application. In the perception layer, intelligent edge nodes are strategically deployed across various detection sites, including agricultural fields, laboratories, and food production facilities, to capture AMR data. Each edge node incorporates an Orange Pi 5B embedded device running a lightweight YOLOv5-based detection model. In parallel, the node regulates the operation of the portable incubator by maintaining key parameters such as temperature, humidity, and carbon dioxide concentration. The transport layer links multiple intelligent edge nodes, facilitating reliable data transmission and inter-node communication. Finally, in the application layer, real-time data analysis enables advanced capabilities such as calculating spatial and temporal trends in AMR and generating alerts based on evaluation outcomes. At the same time, the algorithms are continuously refined with newly acquired data and redeployed to the edge nodes to maintain system adaptability and accuracy.

**Fig. 2** presents a composite view of the web interface of the global AMR monitoring system, which integrates global site visualization with spatial, temporal, compositional, and performance analytics for AMR surveillance across the food supply chain. The central 3D globe depicts spatial risk distributions, while surrounding charts summarize the composition of resistance types, temporal trends across antibiotic classes, system performance indicators, monthly resistant sample counts, and key statistical summaries. Collectively, these components provide an integrated framework to support decision-making and rapid interpretation of global AMR dynamics in the food supply chain. The data currently displayed on the website are primarily illustrative, with real surveillance records available from three pilot sites located in Washington State (USA), British Columbia (Canada), and Berlin (Germany). The platform is designed for scalability, enabling seamless incorporation of AMR datasets from multiple sources through a unified data standard. The development roadmap from pilot testing to a fully operational global surveillance network is outlined in **Fig. S1**. Stage 1 involves completing system design, cloud platform development, and pilot implementation. Stage 2 focuses on automating multi-source data integration, expanding regional surveillance networks, and providing capacity-building through training. Stage 3 targets global deployment, prioritizing high-risk regions and fostering collaboration with international agencies.

**Fig. 2.**
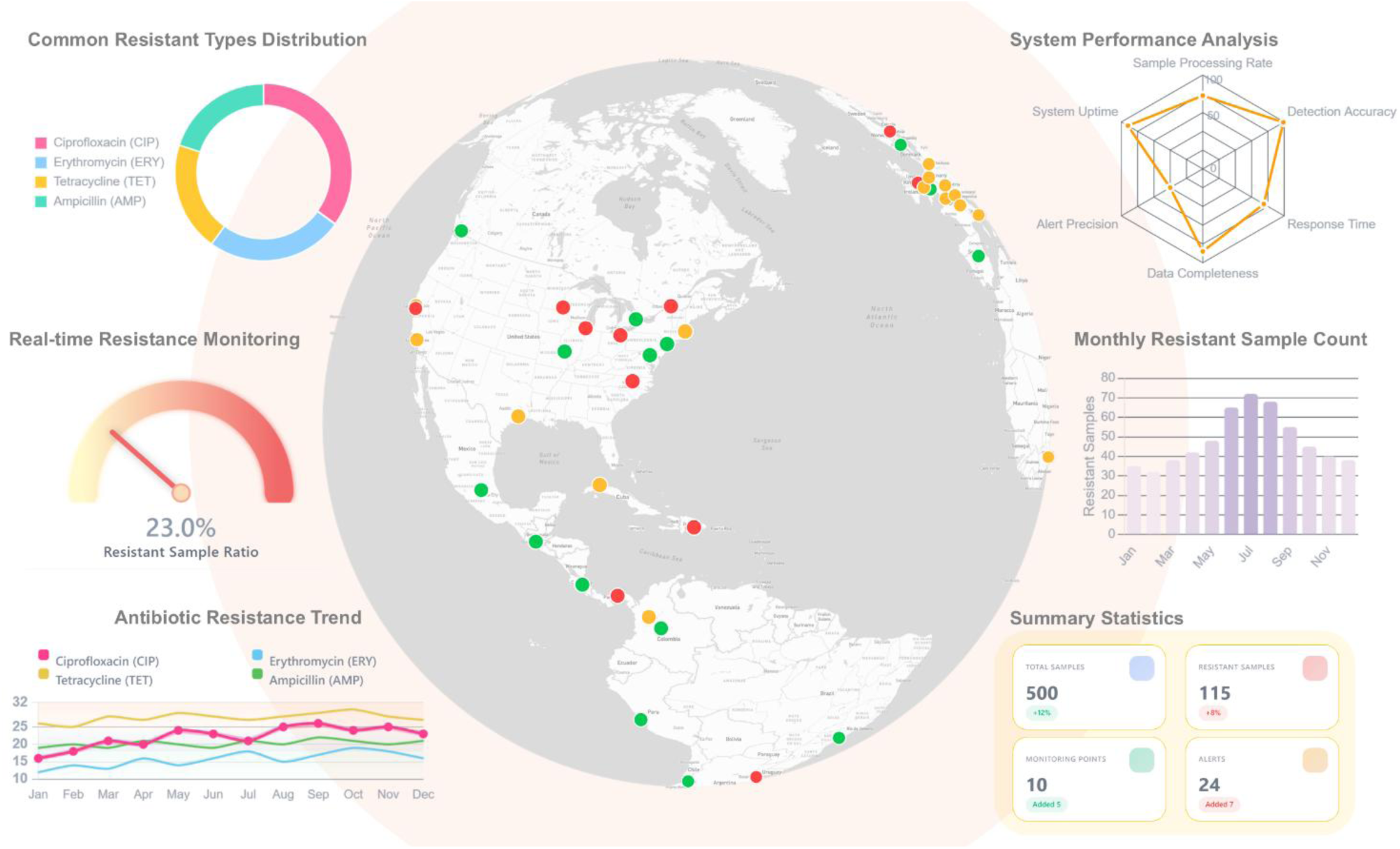
Composite visualization of functional modules within the global AMR online monitoring system for the food supply chain. The central 3D interactive globe displays monitoring sites worldwide and color-coded by AMR risk level (green: low, yellow: moderate, red: high). Real surveillance data are currently available from three pilot locations, namely Washington State (USA), British Columbia (Canada), and Berlin (Germany), with additional points generated for demonstration. Surrounding panels showcase key analytical outputs: a donut chart illustrating the proportion of major resistance types (upper left); a real-time gauge indicating the proportion of resistant samples (middle left); a line chart tracking temporal resistance trends across antibiotic classes (bottom left); a radar chart summarizing system performance metrics (upper right); a bar chart of monthly resistant sample counts (middle right); and summary statistics on total samples, resistant samples, monitoring points, and active alerts (bottom right). Together, these outputs integrate spatial risk mapping, compositional analysis, temporal monitoring, and system performance assessment for global AMR surveillance in the food supply chain.

### 2.2. Portable system for cultivation and monitoring of foodborne pathogens

**Fig. 3a** presents a representative example of a homemade, portable AMR surveillance node, which serves as a key component of the perception layer in the proposed AMR surveillance system. This AIoT-based device powered by an Orange Pi single-board computer (highlighted in the orange box) enables simultaneous testing of pathogen resistance against multiple antibiotics using a portable and low-cost multiplex microfluidic chip as described in the *Methods* section. This fully automated workflow minimizes the need for highly trained personnel, reduces inter-node variability, and supports standardized surveillance across diverse environments, especially in resource-limited settings. Its compact design ensures easy deployment and reliable on-site testing without compromising analytical accuracy.

**Fig. 3.**
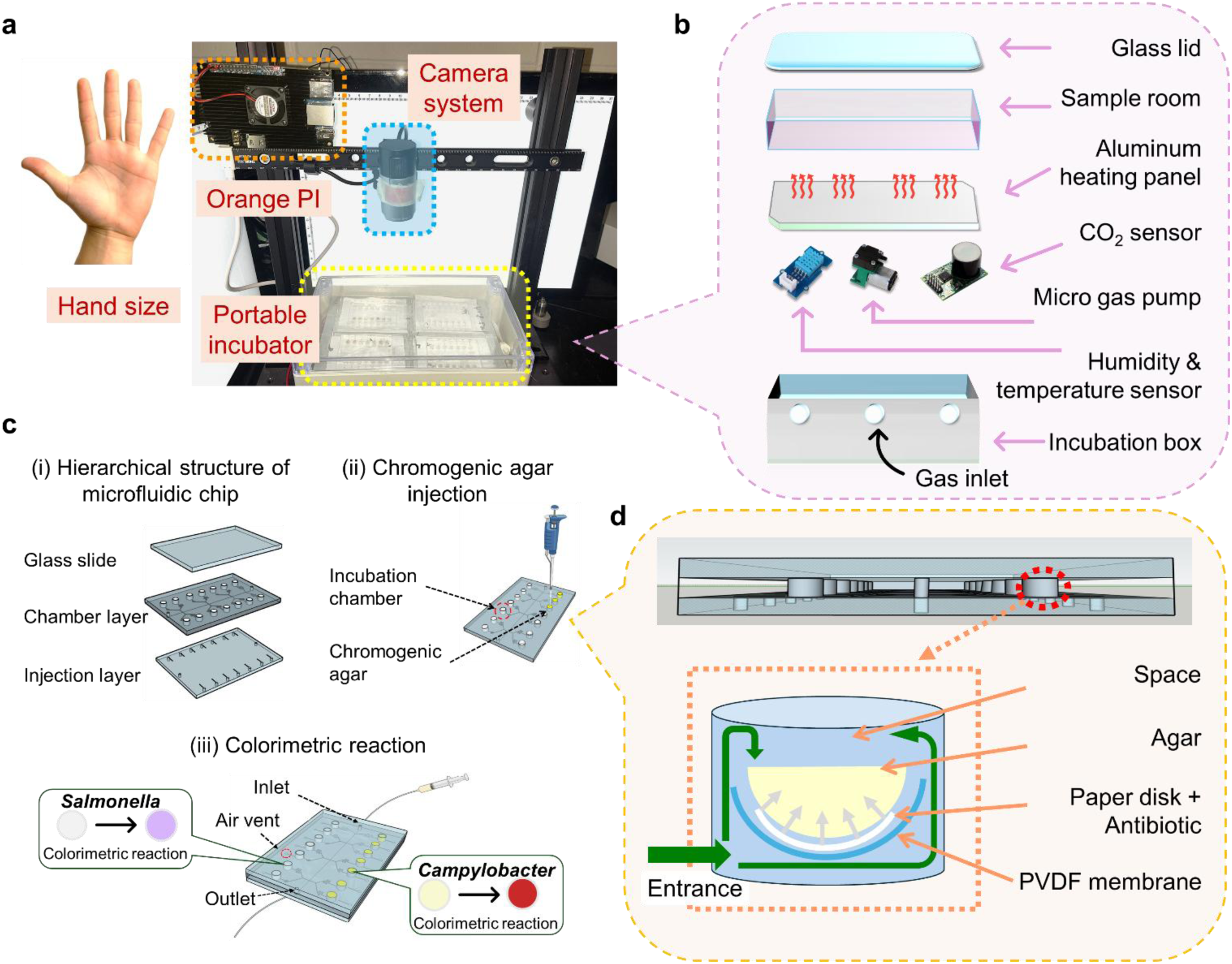
Details of the intelligent edge node. a) Prototype photograph of the AIoT hardware with hand-size comparison. b) Structural layout of the custom-built portable incubator. c) Hierarchical design of the microfluidic chip, showing chromogenic agar injection and colorimetric reaction monitoring. The *Salmonella*-selective agar (left) is off-white by default, whereas the *Campylobacter*-selective agar (right) is light yellow. Growth of antibiotic-resistant bacteria induces a color change in the agar, driven by metabolites released during bacterial growth. d) Design of the incubation chamber within the microfluidic chip. The liquid sample enters through the green inlet and interacts with agar pre-infused with antibiotics.

**Fig. 3b** schematically illustrates the structure of the portable CO₂ incubator integrated into the surveillance node. This incubator plays a central role in AMR monitoring by providing a controlled environment for conducting multiple antibiotic resistance assays in parallel. The incubation chamber contains all essential components, including a micro gas pump, humidity and temperature sensors, and a CO₂ sensor, all managed by the Orange Pi. Sidewall gas inlets allow precise regulation of the incubation atmosphere to promote optimal microbial growth. Above the sensors and pump, an aluminum heating pad equipped with multiple electrodes ensures uniform temperature distribution. A transparent glass lid serves as the chamber ceiling, allowing a 13-megapixel camera mounted above to capture high-resolution images throughout incubation. Sensor data are processed in real time by the Orange Pi, enabling adaptive environmental control and consistent assay performance.

**Fig. 3c(i)** illustrates the structure of the microfluidic chip that consists of a PDMS (polydimethylsiloxane) bottom layer for sample injection, a PDMS middle layer containing 14 incubation chambers, and a top glass layer for sealing and imaging. Chromogenic agar specific to *Campylobacter* or *Salmonella* strains is pre-loaded into these chambers [**Fig. 3c(ii)**] to facilitate bacterial detection via colorimetric reaction [**Fig. 3c(iii)**] in the presence of antibiotic-loaded filter paper (**Fig. 3d**), achieving simultaneous AMR testing against different antibiotics within a single microfluidic chip. **Fig. 3d** schemes a detailed design of the incubation chamber (cross-sectional view) located in the middle layer of the microfluidic chip. During a typical AMR test, the sample liquid is introduced through the inlet port and distributed into each chamber, where a PVDF (polyvinylidene fluoride) membrane supports the chromogenic agar infused with antibiotics diffused from the paper disk. If the bacteria in the sample are resistant to a given antibiotic, they will grow and induce a color change in the agar, which is then captured by the camera for system analysis.

### 2.3. Optimization of on-chip AMR test

#### 2.3.1. Optimization of color response for Campylobacter and Salmonella

*Campylobacter* chromogenic agar changes from yellow to red in the presence of *Campylobacter* species (e.g., *C. jejuni*, *C. coli*, *C. lari*), while *Salmonella* chromogenic agar changes from white to purple in the presence of *Salmonella*. To optimize incubation time, different supplement concentrations were tested for each agar, namely 0.21, 0.42, 0.63, 0.84, 1.05 and 1.26 µg/mL for *Campylobacter* chromogenic agar and 6, 12, 18, 24, 30 and 36 µL/mL for *Salmonella* chromogenic agar. *C. jejuni* F38011 (10^8^ CFU/mL) and *S*. Enteritidis 43353 (10^5^ CFU/mL) were introduced into the microfluidic chip and incubated at 42°C for 60 h with images acquired every 12 h (**Fig. S2a**). For *Campylobacter*, detectable results were achieved within 12 h at a supplement concentration of 0.84 µg/mL. Increasing the concentration to 1.05 or 1.26 µg/mL did not significantly accelerate the color change (**Fig. S2a top**). For *Salmonella*, a minimum concentration of 18 µL/mL was necessary to reduce the incubation time to <24 h (**Fig. S2a bottom**). Therefore, the optimal supplement concentrations were determined to be 0.84 µg/mL for *Campylobacter* and 18 µL/mL for *Salmonella*.

Six bacteria were selected to evaluate the specificity of the chromogenic agars under on-chip incubation conditions. *S.* Enteritidis, *S. aureus*, *L. monocytogenes*, *E. coli*, and *P. aeruginosa* were grown overnight in LB (Luria–Bertani) broth at 37°C under aerobic conditions, while *C. jejuni* was cultivated in MH (Mueller–Hinton) broth under microaerobic conditions at 37°C for 16-18 h. Each bacterial culture was adjusted to 10^8^ CFU/mL before injection into the microfluidic chip, which was then incubated in a portable incubator under microaerobic conditions at 42°C for 60 h. As shown in **Fig. S2b**, color changes occurred only in the presence of the target bacteria (*C. jejuni* and *S.* Enteritidis), confirming the specificity of both agars.

As *Campylobacter* and *Salmonella* are major bacterial pathogens frequently detected together in food products, we further evaluated the sensitivity of the on-chip AMR test for co-cultures of *C. jejuni* F38011 and *S*. Enteritidis 43353 (**Fig. S2c**). Background concentrations were set at 10⁸ CFU/mL for *Campylobacter* and 10⁵ CFU/mL for *Salmonella*, and colorimetric responses were evaluated across inoculation levels of 10², 10⁴, 10⁶, and 10⁸ CFU/mL. On *Campylobacter* chromogenic agar, a red color developed within 12 h at the highest load (10⁸ CFU/mL) with response times increasing as initial concentrations decreased. The detection limit was 10² CFU/mL within 60 h of incubation (**Fig. S2c left**). A comparable pattern was observed for *Salmonella* chromogenic agar (**Fig. S2c right**) although with shorter incubation times, which is consistent with the higher growth rate of *Salmonella* ^26^.

#### 2.3.2. Sensitivity and AST for pathogenic bacteria in food samples

Poultry products are frequently contaminated with both *Campylobacter* and *Salmonella*, making them suitable models for evaluating the performance of the developed microfluidic testing system. Raw chicken breast samples were cut, surface-sterilized, inoculated with defined concentrations of target bacteria, and processed for bacterial recovery as described in the *Methods* section. Four target species were tested in the presence of background organisms: *C. jejuni* F38011 and *C. coli* RM1875 (initial loads: 10², 10⁴, 10⁶, and 10⁸ CFU/25 g) with background *S.* Enteritidis 43353 (10⁵ CFU/25 g); and *S.* Enteritidis 43353 and *S.* Typhimurium S1501 (initial loads: 10², 10⁴, 10⁶, and 10⁸ CFU/25 g) with background *C. jejuni* F38011 (10⁸ CFU/25 g), as shown in **Fig. 4a**. The detection limit for both *Campylobacter* and *Salmonella* was 10⁴ CFU/25 g of chicken. Due to dilution during recovery with phosphate-buffered saline (PBS), the overall turnaround time for food sample testing was longer than for pure cultures with a slight reduction in sensitivity. Importantly, the sensitivity can be further improved with extended incubation, enabling reliable detection of low pathogen concentrations that may occur in food products.

**Fig. 4.**
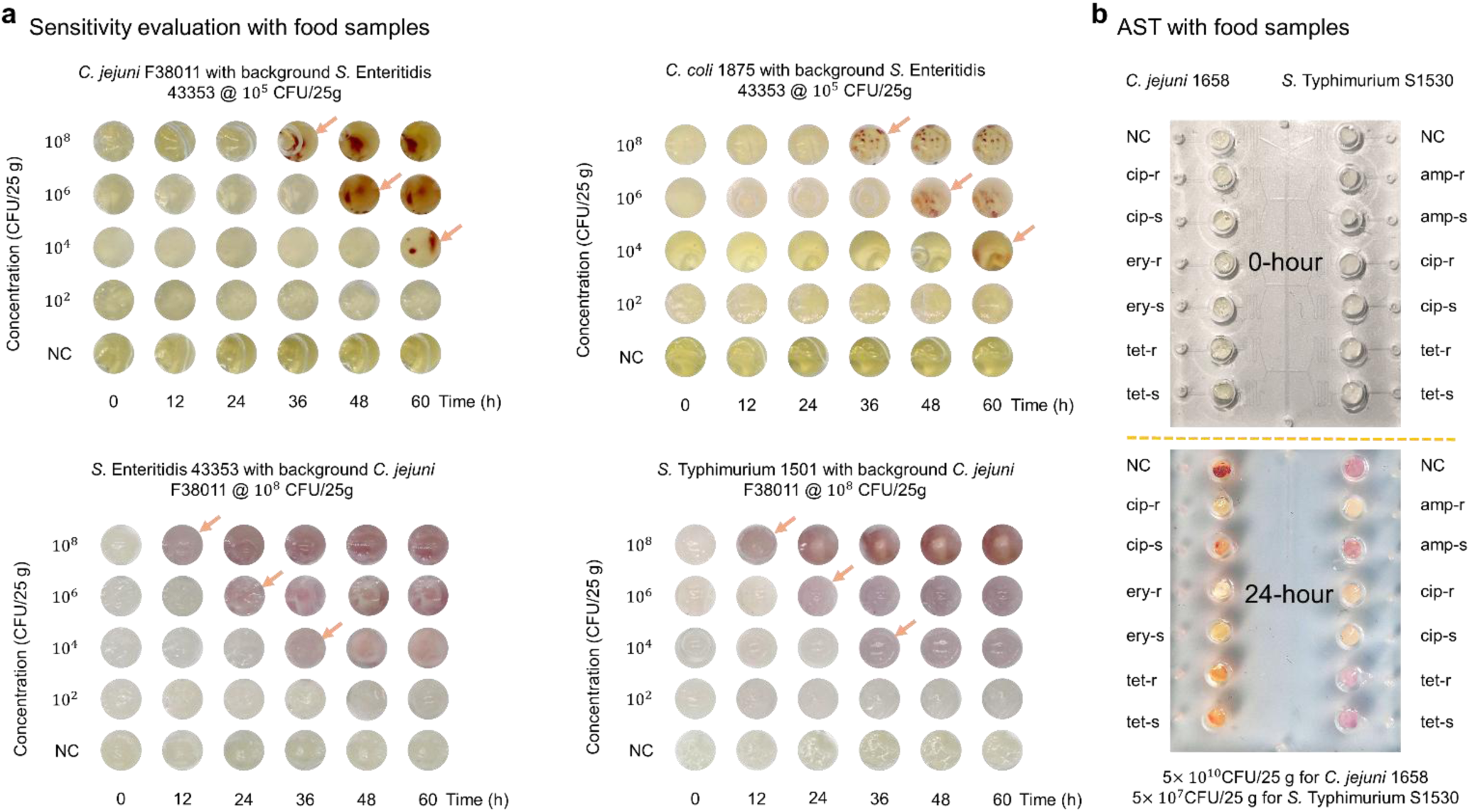
Sensitivity and antimicrobial susceptibility testing (AST) of microfluidic chips for pathogenic bacteria in food samples. a) Sensitivity evaluation of microfluidic chip–based detection of *Campylobacter* and *Salmonella* in chicken meat. Known concentrations of either *C. jejuni* F38011 or *C. coli* RM1875 (10², 10⁴, 10⁶, or 10⁸ CFU/25 g; each with background *S.* Enteritidis 43353 at 10⁵ CFU/25 g), or *S.* Enteritidis 43353 or *S.* Typhimurium S1501 (same concentrations; each with background *C. jejuni* F38011 at 10⁸ CFU/25 g), were spiked onto chicken surfaces to simulate co-contamination. After inoculation, bacteria were recovered with 25 mL phosphate-buffered saline (PBS), applied to microfluidic chips, and incubated at 42°C for 60 h. Arrows indicate visible color changes corresponding to positive detection. b) Representative on-chip AST results for *C. jejuni* 1658 (left) and *S.* Typhimurium S1530 (right) recovered from spiked chicken samples after 24 h of cultivation. Initial inoculation levels were 5 × 10¹⁰ CFU/25 g for *C. jejuni* 1658 and 5 × 10⁷ CFU/25 g for *S.* Typhimurium S1530. Each microfluidic chamber contained one of the following: ciprofloxacin (cip), erythromycin (ery), tetracycline (tet), or ampicillin (amp) at resistant (-r) or susceptible (-s) breakpoint concentrations, or no antibiotics (negative control, NC). Images show chips at 0 h and after 24 h of incubation. For *C. jejuni*, growth (red) was observed in NC, cip-s, tet-r, and tet-s chambers. For *S.* Typhimurium, growth (purple) appeared in NC, amp-s, tet-r, and tet-s chambers, indicating resistance to these antibiotics.

For AST with chicken samples, higher initial bacterial loads were required to ensure sufficient recovery for standard protocols, yielding an overall recovery rate of ∼10%. *C. jejuni* 1658 and *S.* Typhimurium S1530 were selected for AST tests. Chicken samples were inoculated with 5 × 10¹⁰ CFU/25 g of *C. jejuni* 1658 and 5 × 10⁷ CFU/25 g of *S.* Typhimurium S1530. The recovered bacteria were then subjected to on-chip AST. Specifically, *C. jejuni* 1658 was tested against ciprofloxacin (CIP), erythromycin (ERY), and tetracycline (TET), while *S.* Typhimurium S1530 was tested against ciprofloxacin (CIP), ampicillin (AMP), and tetracycline (TET). For each antibiotic, both resistant (-R) and susceptible (-S) concentrations were applied according to the CLSI guidelines ^27^ as summarized in **Table S1**, along with a negative control (NC) well containing no antibiotic. As shown in **Fig. 4b**, colorimetric changes were compared between 0 h and 24 h. For *C. jejuni* 1658, growth (red signal) was observed in NC, CIP-R, TET-R, and AMP-R chambers. For *S.* Typhimurium S1530, growth (purple signal) appeared in NC, CIP-R, AMP-R, and TET-R chambers. These results were consistent with MIC testing, confirming resistance to the respective antibiotics.

### 2.4. AIoT nodes designed for AMR detection

The AIoT-enhanced node was designed and implemented to automate the recognition of chromogenic reactions in microfluidic chips, improving AMR surveillance and data collection efficiency. **Fig. 5** demonstrates the workflow of Orange Pi and outlines the creation of an intelligent edge node that comprises three key components, namely original data collection and annotation (**Fig. 5a**), model fine-tuning (**Fig. 5b**), and model deployment (**Fig. 5c**). In the data collection and annotation module, a high-resolution IoT camera captured images of microfluidic chips during bacterial cultivation, recording the growth and associated colorimetric changes. These images were processed and annotated using advanced computer-vision tools, producing a high-quality dataset for model training. In the fine-tuning stage, the annotated dataset was used to customize the YOLOv5 model, enabling accurate detection of microfluidic wells not included in the original pre-set categories, as well as reliable identification of chromogenic changes associated with AMR *Salmonella* or *Campylobacter*. The deployment stage involved model compression to reduce computational load with minimal loss of accuracy, followed by cross-compilation to ensure compatibility between standard ×86_64 architectures and the ARM-based Orange Pi 5B. The optimized model operated in real time on the Orange Pi device, successfully monitoring colorimetric reactions indicative of AMR profiles for *Salmonella* and *Campylobacter*.

**Fig. 5.**
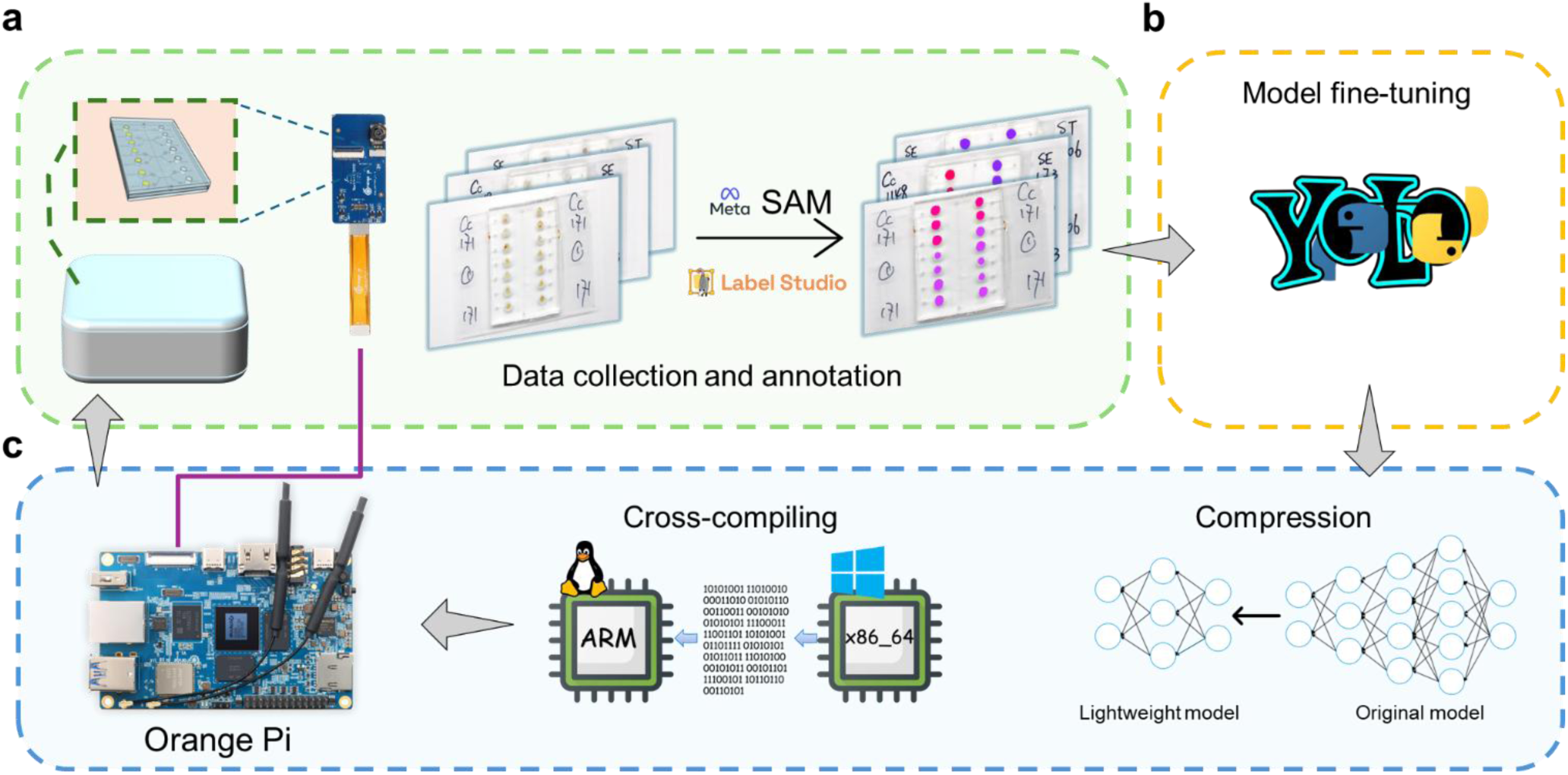
Construction workflow of intelligent edge nodes for automated microfluidic chip analysis. The workflow comprises three major components: a) Data collection and annotation. Orange Pi automates bacterial cultivation and captures images from microfluidic chips. Image annotation is conducted in Label Studio, assisted by the Segment Anything Model (SAM), a large-scale computer vision framework, to generate high-quality training datasets. b) Model fine-tuning. Annotated data are used to fine-tune a YOLOv5 model for object detection and classification. c) Model deployment. The optimized model is compressed into a lightweight version, cross-compiled for the ARM architecture, and redeployed onto the Orange Pi for efficient edge inference and real-time processing.

#### 2.4.1. Data collection and annotation

In total, 219 high-resolution images of microfluidic chips were selected, each containing one to four chips with 14 wells per microfluidic chip and capturing various stages of pathogen growth along with their corresponding colorimetric reactions. Data augmentation (grayscale conversion, noise addition, cropping, rotation) expanded the dataset to over 1,000 images comprising more than 15,000 annotated well objects, ensuring sufficient diversity for robust model training ^28^. Accurate dataset annotation is critical for guiding machine learning models to identify object locations and categories within images. In this study, the labeling process was significantly accelerated using SAM (Segment Anything Model) ^29^, which is a foundational computer vision model developed by Meta and trained on 1.1 billion masks for annotation. Combined with the front-end interface of an open-source data annotation tool Label Studio, this approach enabled rapid and precise labeling of all wells in the microfluidic chips, including their borders and bounding boxes. Compared with manual annotation, this strategy greatly reduced labeling time while maintaining high accuracy. Consequently, large-scale images were annotated efficiently, yielding a high-quality dataset for subsequent model fine-tuning. Representative annotation results are shown in **Fig. S8**.

#### 2.4.2. Fine-tuned YOLOv5 model performance

The fine-tuned YOLOv5 model demonstrated excellent performance in detecting wells and identifying color changes in microfluidic chip images. By leveraging data augmentation and optimized training strategies, the model achieved over 99.5% accuracy across three key metrics, namely Precision, Recall, and mAP. Manual identifications were used as the ground truth for validation, and the model showed strong concordance with human evaluation in correlating colorimetric changes with bacterial AMR patterns. A training log, including loss convergence and metric evolution, is presented in **Fig. S15** and highlights rapid convergence and stable performance across long epochs. Collectively, these results confirm that the customized YOLOv5 model can robustly and accurately identify well positions and chromogenic reaction outcomes in the microfluidic chips.

#### 2.4.3. Practical deployment on Orange Pi 5B

Application of a PTQ compression algorithm reduced the YOLOv5 model size by ∼80% while preserving nearly identical accuracy. When deployed on the Orange Pi 5B (ARM64), the optimized model achieved an inference speed of ∼60 FPS, a 1.5× improvement over the baseline Float32 model. Notably, the deployment of ARM64 maintained accuracy consistent with the ×86 baseline, confirming that the compressed model can be executed efficiently on resource-constrained ARM platforms without compromising performance.

### 2.5. Validation of the AIoT-based AST using local bacterial isolates

To assess the reliability of our AIoT-based AMR surveillance platform, we compared its performance with conventional AST methods using local bacterial isolates. While the AI model achieved over 99.5% accuracy in interpreting colorimetric reactions, overall system performance remained dependent on the initial chromogenic signal generated within the microfluidic chip. Therefore, benchmarking against traditional AST methods as a well-established standard for susceptibility assessment was essential to validate practical applicability beyond theoretical accuracy.

We evaluated the platform using two antibiotic-susceptible reference strains (*C. jejuni* F38011 and *S.* Enteritidis 43353), along with six potentially resistant *Campylobacter* isolates and six *Salmonella* isolates (**Fig. 6a, b**). Repeated batch testing was performed for all strains listed in **Table S2**. The final categorical agreement rates between the AIoT system and conventional AST were 96.76% for *Salmonella* and 95% for *Campylobacter* (**Table S3**), both exceeding the U.S. Food and Drug Administration’s minimum threshold of 90% categorical agreement. These findings demonstrate the robust performance and practical potential of the AIoT-based AST platform compared with the standard methodologies.

**Fig. 6.**
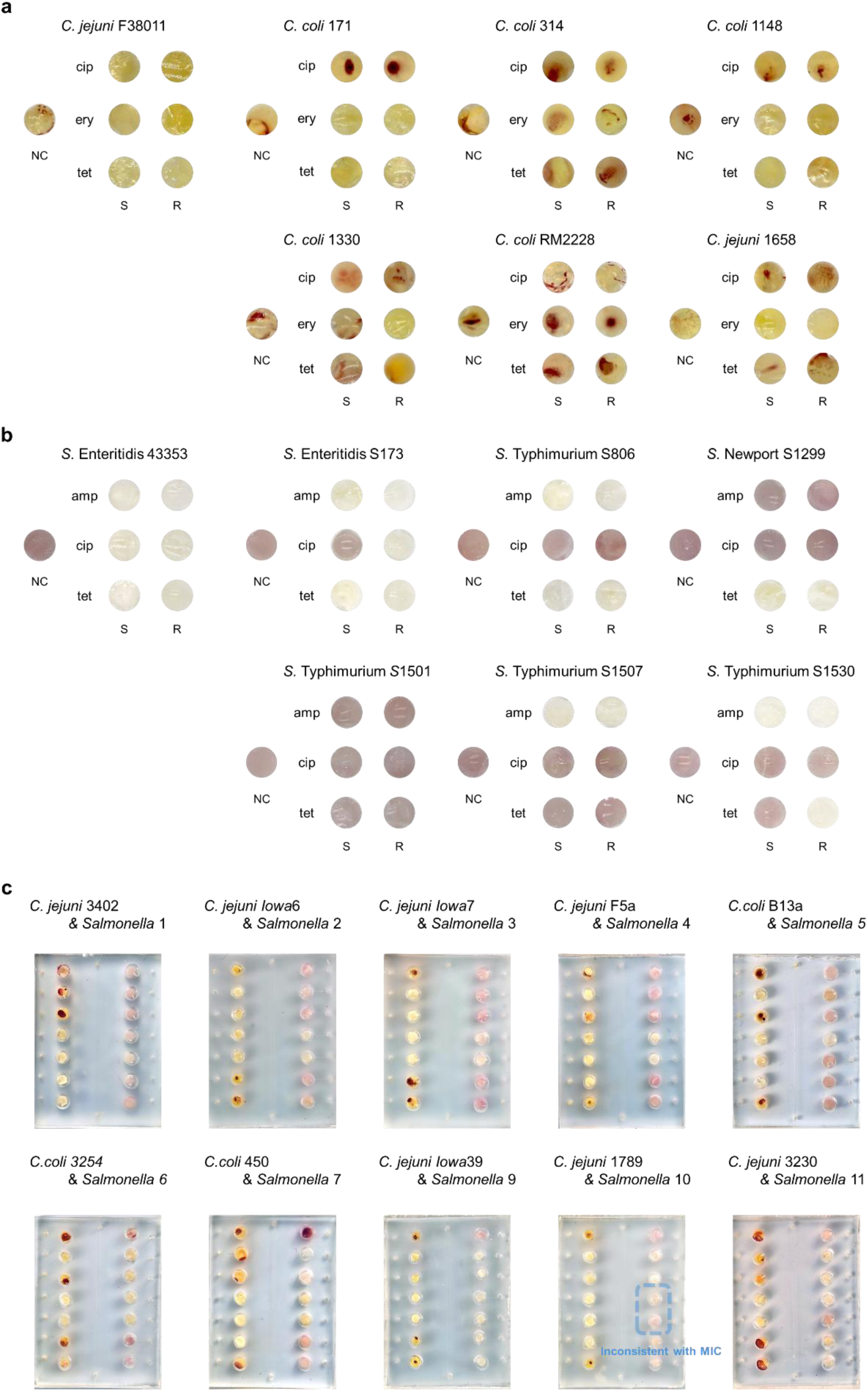
On-chip antimicrobial susceptibility testing (AST) validation for determining multidrug resistance profiles of locally and internationally isolated *Campylobacter* and *Salmonella* strains. Each chamber of the microfluidic device was loaded with one antibiotic, namely ciprofloxacin (cip), erythromycin (ery), tetracycline (tet), or ampicillin (amp), at susceptible (-S) or resistant (-R) breakpoint concentrations; chambers containing agar without antibiotics served as negative controls (NC). a, b) Multidrug resistance profiles of seven locally isolated *Campylobacter* strains (*C. jejuni* F38011, *C. coli* 171, *C. coli* 314, *C. coli* 1148, *C. coli* 1330, *C. jejuni* 1658, and *C. coli* RM2228) and seven locally isolated *Salmonella* strains (*S.* Enteritidis 43353, *S.* Enteritidis S173, *S.* Typhimurium S806, *S.* Newport S1299, *S.* Typhimurium S1501, *S.* Typhimurium S1507, and *S.* Typhimurium S1530) determined using the microfluidic platform. c) On-chip validation results of internationally isolated *Campylobacter* (left) and *Salmonella* (right). In the *Campylobacter* columns, the first chamber is NC (negative control), followed from top to bottom by cip, ery, and tet at resistant (-R) or susceptible (-S) breakpoint concentrations. In the *Salmonella* columns, the first chamber is NC, followed from top to bottom by ampicillin, ciprofloxacin, and tetracycline at resistant (-R) or susceptible (-S) concentrations. A red signal in the *Campylobacter* chamber or a purple signal in the *Salmonella* chamber indicates bacterial growth at the corresponding antibiotic concentration.

### 2.6. International validation of the AIoT-based AST platform using multi-regional bacterial isolates

To further evaluate the robustness of the AIoT-based AST platform across diverse contexts, we conducted international validation using *Salmonella* and *Campylobacter* isolates collected from different geographic regions (USA, Canada and Germany). Building on prior validation with local field isolates, this further evaluation assessed the consistency and accuracy of the platform across a broader range of genetic backgrounds and antimicrobial resistance lineages. In total, 10 *Salmonella* and 10 *Campylobacter* isolates were tested and benchmarked against conventional AST methods. The categorical agreement rates were 96.76% for *Salmonella* and 100% for *Campylobacter*, closely aligning with results from local isolates and exceeding the FDA’s minimum requirement of 90% categorical agreement. These findings confirm that the platform delivers high accuracy and reliability even when challenged with genetically diverse isolates at an international scale. The on-chip AST results are shown in **Fig. 6c** with detailed isolate information provided in **Table S4**.

## 3. Discussion

This study presents the development of an automated AMR monitoring system that integrates microfluidic technology with AIoT components, specifically tailored for application in the food supply chain. This platform combines miniaturized on-chip testing, advanced computer vision algorithms, and distributed AIoT infrastructure to address key limitations of conventional AMR surveillance in a cost-effective and scalable manner. Compared with existing AST systems, our approach introduces several notable improvements. First, the integration of colorimetric microfluidic chips with on-device deep learning enables automated, accurate, and rapid determination of resistance phenotypes. This system achieved categorical agreement rates exceeding 95% across both local and multi-regional bacterial isolates. This accuracy is comparable to conventional gold standard broth microdilution while additionally offering high-throughput batch testing and substantial cost savings, reducing expenses by up to 98% (**Table S5**). Second, by incorporating AIoT nodes with real-time environmental sensing and cloud connectivity, this platform supports decentralized testing and immediate data aggregation. This design directly addresses persistent challenges of cost, labor intensity, and reporting delays, particularly in resource-limited settings. The real-time and scalable nature of this system has important public health implications. Unlike national and international AMR surveillance programs that often rely on centralized and retrospective data analysis, our approach enables daily resistance monitoring across multiple sites. This higher temporal and spatial resolution is especially valuable within global food production and distribution networks, where resistant strains can spread rapidly across borders. Nonetheless, several limitations should be acknowledged. The current validation has been restricted to *Salmonella* and *Campylobacter*. Future work should broaden the coverage of the platform to include a wider range of foodborne and environmental pathogens. In addition, variations in environmental factors, such as lighting conditions or sample composition, could also influence AI accuracy, underscoring the need for continued refinement of detection algorithms. For future works, expanding the spectrum of antibiotics and bacterial species tested will improve generalizability. Integrating the system with global AMR databases could facilitate international data sharing and provide actionable insights for policy development. Pilot studies conducted across the food supply chain including farms, processing facilities, and retail environments will be essential to evaluate the practicality, scalability, and overall impact of this platform. In conclusion, this work demonstrates the feasibility and advantages of next-generation AIoT-enabled AMR surveillance within the food supply chain. By delivering rapid, automated, and scalable resistance profiling at the point of need, this platform has the potential to transform global AMR monitoring strategies and contribute to enhanced food safety outcomes.

## 4. Methods

### 4.1. Bacterial strains, cultivation conditions, and antibiotics

Eight *Campylobacter* strains and seven *Salmonella* strains from different sources were selected for on-chip testing (**Table S2**). Four other bacteria were used for the specificity test, including *S. aureus* MRSA-10, *Listeria monocytogenes* ATCC 7644, *E. coli* K12, and *P. aeruginosa* PA14. These strains were selected to represent phylogenetically diverse microorganisms relevant to the global food supply chain, including common foodborne pathogens and frequent environmental or commensal microbes. These bacterial stains were cultivated in Luria-Bertani (LB) broth in aerobic conditions at 37°C overnight. All *Campylobacter* strains were routinely cultivated on Mueller-Hinton (MH) agar (BD Difco; Fisher Scientific, Canada) supplemented with 5% (v/v) defibrinated sheep blood (Abbott, Canada) and incubated in the microaerobic conditions (85% N_2_, 10% CO_2_, 5% O_2_) at 37°C for 48 h. Overnight *Campylobacter* culture was prepared by transferring bacterial colonies from the agar plates to MH broth (BD Difco; Fisher Scientific, Canada), followed by incubation in the microaerobic conditions at 37°C for 16-18 h with constant shaking at 175 rpm. All *Salmonella* strains were grown on LB agar in the aerobic conditions at 37°C for 24 h. Overnight *Salmonella* culture was prepared by transferring bacterial colonies from the agar plate to LB broth (BD Difco; VWR, Canada), followed by incubation in the aerobic conditions at 37°C for 12 h with constant shaking at 175 rpm.

Ampicillin sodium salt, ciprofloxacin, erythromycin, and tetracycline hydrochloride (Sigma-Aldrich, Canada) were selected as representative antibiotics for the AMR tests, as they are widely used in the clinical and veterinary treatment of *Salmonella* and *Campylobacter* infections and serve as key markers in global AMR surveillance due to rising resistance trends in these genera. The stock solutions (640 µg/mL) were prepared by dissolving ampicillin, erythromycin, and tetracycline in distilled water and ciprofloxacin in dimethyl sulfoxide, followed by passing through a sterile nylon syringe filter (0.22-µm pore size) and stored at -20°C. Upon on-chip testing, the stock solutions were freshly diluted with sterile distilled water to the desired concentrations, and a 2-µL aliquot of each solution was applied to each sterile paper disk (2 mm in diameter, Whatman No. 1 filter paper; Fisher Scientific, Canada). The paper disks were air-dried in a biosafety cabinet before being placed into the incubation chambers of the microfluidic chip.

### 4.2. Design and fabrication of the microfluidic chip

The microfluidic chip consists of three layers. The top layer is a plain glass microscope slide (75 × 50 × 1 mm; Fisher Scientific, Canada), while the bottom and middle layers are polydimethylsiloxane (PDMS) slabs with designed microfluidic patterns. The top glass layer provides a clear window for image data collection and seals the middle layer. The middle layer with microchambers serves as the incubator, and the bottom layer provides the inlets for sample injections and function as the base seal for the entire chip. The PDMS slabs for the bottom and middle layers were fabricated following the typical protocol with modifications ^30^, including the design of microfluidic patterns, preparation of a master by photolithography, and preparation of PDMS slabs using the master mold.

The microfluidic patterns for the middle and injection layers were designed and printed as photomasks by CAD/Art Services (Bandon, OR, USA). The middle layer includes an inlet port (⌀=1.5 mm) and an outlet port (⌀=1.5 mm) connected by a main channel with a width of 500 µm. On each side of this main channel, 14 incubation chambers (7 on each side, ⌀=4 mm) are connected to the channel by a serpentine microchannel (width=100 µm). Each incubation chamber is further connected to a positioning disk (⌀=1.5 mm) to facilitate alignment with the venting ports in the bottom layer (**Fig. S3a**). The bottom layer has inlet and outlet ports aligned with those in the middle layer (**Fig. S3b**). It also includes 14 positioning disks (⌀=4 mm) corresponding to the incubation chambers in the middle layer, with each connected to a venting port (total 14; ⌀=1.5 mm) via a microchannel (width = 100 µm) for exporting excess fluids.

The protocol of photolithography was adapted to the equipment in the cleanroom at the McGill Nanotools Micro and Nanofabrication Facility. First, 5 mL of SU-8 2050 photoresist (Kayaku Advanced Materials Inc., USA) was spin-coated onto a clean silicon wafer (⌀ = 100 mm, University Wafer, USA). The spin coating was performed at 500 rpm for 5 s, followed by 2000 rpm (acceleration rate = 1305 rpm/s) for 45 s to achieve an even layer with thickness of 80 µm corresponding to the height of the microfluidic channel. Then, the wafer underwent soft baking at 65°C for 3 min, followed by 95°C for 9 min, and allowed to cool to room temperature. Next, the wafer with the cured SU-8 was covered by the photomask and exposed to UV light with a dose of 215 mJ/cm^2^, followed by a post-exposure baking at 65°C for 2 min and 95°C for 7 min. After baking, the wafer was developed in SU-8 developing solvent (Kayaku Advanced Materials Inc., USA) with occasional shaking until the unexposed SU-8 was fully removed, and excessive solvent was rinsed off and dried with nitrogen gas. Last, the wafer with the developed patterns underwent hard baking in a vacuum oven at 200°C for 5 min and was allowed to cool to room temperature, which was ready to serve as the master mold.

To fabricate the PDMS slabs, the precursor and curing agent (Dow Sylgard 184 silicone encapsulant clear kit, Canada) were mixed at a weight ratio of 10:1 and poured onto the master mold with an aluminum disc (18 g for the bottom layer and 30 g for the middle layer). After degassing in a vacuum chamber, the disc was heated on a hot plate at 80°C for 20 min to accelerate the curing process. The obtained elastomer slabs were then peeled from the molds, cut to rectangular shapes, and punched using Miltex biopsy punches (Ted Pella Inc., USA) at the locations of the chambers (middle layer), inlet/outlet (middle and bottom layers), and venting ports (bottom layer).

The assembly of the microfluidic chip started with binding the two PDMS slabs together by plasma treatment and heating. After that, a single autoclaved PVDF membrane (⌀ = 4 mm) was placed into each incubation chamber, followed by placement of an antibiotic-loaded paper disk (⌀ = 2 mm). Next, 20 µL of either *Campylobacter* chromogenic agar medium (CHROMagar Campylobacter; CHROMagar, USA) or *Salmonella* chromogenic agar medium (CHROMagar Salmonella Plus; CHROMagar, USA) was individually loaded into each incubation chamber. Last, the plain glass slide was used to seal the PDMS layers from the top by plasma treatment to complete the assembly of the microfluidic chip (**Fig. S4**). The entire microfluidic chip was then sterilized under UV light for 30 min and sealed for storage.

After assembling the microfluidic chip, a cross-contamination test among the incubation chambers was conducted (**Fig. S5**). Paper disks with or without blue dye were placed into the incubation chambers of the chip (**Fig. S5a**). A yellow dye solution was then injected to fill all the chambers completely. The chambers that initially contained blue dye turned green due to mixing with yellow dye, while other chambers remained yellow (**Fig. S5b**). To assess the potential diffusion of the green dye, the microfluidic chip was placed in a CO_2_ incubator for 60 h to simulate bacterial cultivation and detection conditions. Chambers containing only yellow dye showed no color change, indicating no detectable diffusion of the green dye occurred from adjacent chambers (**Fig. S5c**). This result confirms that the microfluidic chip design effectively prevents cross-contamination.

### 4.3. AMR test using microfluidic chips

For each test, the sample fluid (i.e., bacteria culture) was injected into the chip using a syringe pump connected to sterile accessories (i.e., syringe, polyvinyl chloride tubing, elbow adaptors) (**Fig. S6**). All components were stored in sealed conditions and sterilized by either UV light or 70% isopropanol before use or obtained freshly from the sterile pack. A petri dish was used for collecting waste solutions, and a binder clip was placed on the outlet tubing to act as the stopper. The bacterial culture in the syringe was pumped at a rate of 50 µL/min. Once the main channel was filled completely, the binder clip was put on to block the outlet tubing, forcing the liquid to flow into the chambers on both sides. After all chambers were fully filled with sample fluid (20 µL), the pump was stopped, and clear tape was used to seal the inlet, outlet and all the venting ports to avoid leakage and evaporation. Finally, the microfluidic chip was incubated in a homemade incubator (10% CO_2_) at 42°C for up to 60 h.

### 4.4. Preparation and processing of poultry samples for on-chip testing

Raw, boneless, and skinless chicken breast samples were purchased from local supermarkets and processed following the U.S. FDA *Bacteriological Analytical Manual* (BAM) protocol ^31,32^. Using sterile instruments, the chicken breast was cut into 25-g portions. To eliminate native microorganisms and ensure controlled experimental conditions, each portion was soaked in 1% (v/v) sodium hypochlorite solution (Sigma-Aldrich, USA) for surface sterilization. The samples were then inoculated with defined concentrations of *Campylobacter* and *Salmonella* on the surface and subsequently air-dried in a biosafety cabinet. For bacterial recovery, each inoculated sample was placed in a sterile stomacher bag with 25 mL of sterile phosphate-buffered saline (PBS). The bag was hand-massaged for 3 min to release bacteria from the meat surface. The rinse solution was then collected and passed through sterile Whatman No. 1 filter paper to remove tissue debris. The filtrate was used directly for microfluidic chip detection and AST.

### 4.5. Structural features of Orange Pi 5B

Orange Pi is the core of the AIoT system and can run lightweight AI models using its built-in neural processing unit (NPU) ^33,34^. To fully understand how Orange Pi drives the automated detection system, it is essential to explore its hardware design and the corresponding software support (**Fig. S7**). **Fig. S7a** presents a photograph of Orange Pi 5B hardware. Despite its compact and palm-sized design, this device functions as a fully featured computer. It integrates a CPU (central processing unit) for general computing and complex task scheduling, a GPU (graphics processing unit) for graphic-related tasks including video and gaming, and a NPU for accelerating deep learning tasks. These three units are housed on a single chip substrate shown in yellow. The two components labeled in green are storage devices. One is random access memory (RAM) that exchanges data directly with the CPU at high speed but loses all stored data when powered off. The other is eMMC Flash that serves as disk storage, offering larger space for long-term use. The Ethernet port along with Wi-Fi and Bluetooth capabilities is marked in red and enables Orange Pi to communicate with other IoT devices or cloud servers. Orange Pi features numerous input/output (I/O) interfaces with three components highlighted in orange. These include a camera interface for connecting IoT camera to take photos, an HDMI output for monitoring linkage, and a 26-pin general-purpose I/O (GPIO) interface commonly used in electronic devices such as microcontrollers and microprocessors ^35^. The term “26-pin GPIO” refers to an interface that comprises 26 independent pins, each individually programmable as either input or output to serve special functions such as power supply (5V and 3.3V), serial communication (I2C, SPI, UART) ^36^, pulse width modulation (PWM), and analog input. These versatile pins on Orange Pi facilitate connections with various external devices such as LEDs and sensors and enable communication with other microcontrollers ^37,38^.

**Fig. S7b** illustrates the system architecture based on the hardware structure, including a System on a Chip (SoC), memory, external devices, and network within the hardware layer. This system is built around RK3588, which is Rockchip’s latest flagship AIoT SoC with an 8-nm lithography process. It features an 8-core 64-bit ARM architecture CPU with frequencies up to 2.4 GHz, a Mali-G610 quad-core GPU, and a built-in AI accelerator NPU that provides 6 Tops of computing power while supporting mainstream deep learning frameworks. The RK3588S optimizes performance across various AI applications. The SoC communicates data via a high-speed bus connection to memory and uses GPIO for interaction with external devices such as temperature sensors, humidity sensors, heaters, and CO_2_ micro-gas pumps. This system incorporates three methods for remote resource communication, namely Ethernet, WLAN (including Wi-Fi) and Bluetooth. Ethernet and WLAN are ideal for large-scale, high-speed data transmission, while Bluetooth is used for small data exchange or peripheral connection (e.g., keyboard, mouse). In this case, WLAN is primarily used to transmit substantial amounts of data including high-resolution images from the IoT camera and continuous incubator log information (e.g., temperature, CO_2_ concentration) during incubation.

The AIoT device relies on robust software support to function effectively. We selected an open-source Linux system Ubuntu 22.04 LTS that facilitates two primary objectives, namely running the AI model and controlling the hardware. This AI component supports frameworks such as PyTorch ^39^ and TensorFlow ^40^, simplifying the development and execution of deep learning models. The control function primarily comprises custom code used to read data from either built-in or externally connected hardware on Orange Pi and adjust strategies as needed. For example, it continuously monitors CO_2_ level and operates a micro-gas pump to maintain balance. Therefore, Orange Pi satisfies the requirements for AMR surveillance.

### 4.6. Developing a labeling workflow for microfluidic image segmentation and dataset preparation

The labeling system used in this study consists of two main components, namely the Segment Anything Model (SAM) ^29^ and the Label Studio. Powered by Meta AI, SAM provides automated preliminary segmentation through an interactive method for efficient labeling. Label Studio is a widely used open-source data labeling platform that offers a comprehensive toolchain for tasks ranging from data loading to outputting labelled results. It supports various machine learning models for quickly creating image label masks. In this study, SAM was used as the machine learning backend for Label Studio. Both Label Studio v1.2 and the SAM ML backend were deployed by Docker ^41^, which simplifies system requirements and streamlines the installation and maintenance process. Docker was run on a Windows 11 operating system with the following hardware: an Intel 13900K CPU, 128 GB of RAM, and an NVIDIA RTX 4090 GPU. The CUDA toolkit (v12.4) ^42^ and cuDNN library (v8.9.4) ^43^ for GPU acceleration were automatically configured when Label Studio was installed via Docker.

The detailed process of constructing the labeling system is outlined as follows. First, we installed Docker and Git and then cloned the Label Studio ML backend Git repository. Afterward, we constructed the SAM ML backend for our system and then launched the Label Studio via Docker, making it accessible via a web browser at localhost (https://localhost:8080). After registering an account and logging in, we created a new labeling project. An access token was then generated and used to establish a connection between the frontend labelling interface and the SAM ML backend, which supports both key points and bounding boxes labeling assistance. SAM can interactively enhance segmentation accuracy based on user-specified target and non-target key points. We focused on two types of targets (positive and negative holes) and one non-target object (background). During the labeling process, we selected a well in the microfluidic chip, prompting the SAM ML backend to automatically identify similar targets. When the SAM results were not ideal, we refined them by adding key points to the background or targets to guide the model. The target labeling interface and export results are shown in **Fig. S8**.

### 4.7. Architecture of the lightweight YOLOv5s model

YOLO was originally developed on a deep learning framework named Darknet ^44^. YOLO v5 is the first version of YOLO models written on the PyTorch framework, making it easier to use. For setting up YOLO dependencies, we used YOLO v5s release v7.0 as the reference for fine-tuning, with the backbone part frozen. YOLO requires a GPU to operate on PyTorch. We selected recent versions of these dependencies: PyTorch 2.1.2, TorchVison 0.16.2, CUDA 12.1 for an Ubuntu 22.04 LTS system.

The detailed model structure of YOLO v5 is shown in **Fig. S9** and consists of four main components, namely input, backbone, neck, and output. The input to the model is the original image collected by a camera, which is then reshaped into a 640 × 640-pixel format with three color channels (red, green, blue). This 3D tensor serves as the input for the YOLO network. Following the input is the backbone, which is responsible for feature extraction. YOLO v5 utilizes CSPDarknet as its backbone, a combination of CSP (Cross-Stage Partial) and Darknet53, known for enhancing the learning speed and accuracy of CNN. Key components in the backbone include Focus Layer, which reduces layers, parameters, calculation complexity and GPU memory while increasing forward and backward speed with minimal impact on precision. Detailed structure of these functional layers can be found in **Fig. S10-S13**.

Following the backbone is the “neck” of YOLO called PANet (Path Aggregation Network) ^45^, a design methodology for feature fusion and selection to improve object detection performance. PANet enhances information flow by allowing deeper features (closer to the output) to propagate back to shallower features (closer to the input). The backbone and the neck structure (e.g., PANet) serve different purposes. The backbone is responsible for feature extraction from raw images, while PANet or other similar structure is designed to better use those extracted features for prediction tasks. Additionally, backbone is generally pre-trained on a large dataset such as ImageNet ^46^ for image classification, exploiting the transferability of learned features. In contrast, neck structures are usually trained from scratch for specific tasks, such as object detection and segmentation by using pre-extracted features from the backbone.

Following the neck is the “head” of YOLO that primarily serves to convert feature maps into the final output target, such as classes and bounding boxes. YOLO v5 makes predictions on different feature map scales corresponding to different sizes of objects. It typically uses three scales for prediction, one for large objects and two for small ones, represented as three blue cubes in the head shown in **Fig. S9**. For each scale, the model outputs a series of vectors containing category and location information, where each vector corresponds to several bounding boxes tied to a grid of the input image. The head structure varies slightly depending on the task. For example, in an object detection scenario with 2 bounding boxes per grid and 3 categories labeled as *a*, *b*, *c*, the output for each grid can be expressed as Formula (1):

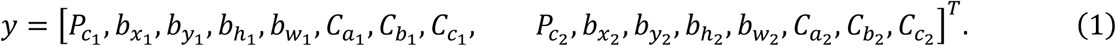

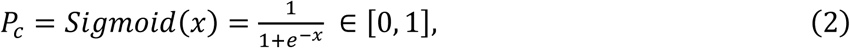

and 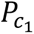 refers to the probability of an object present in the bounding box No.1 within this grid, calculated using a Sigmoid function to constrain its value between 0 and 1. 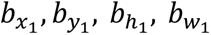 denote the x and y coordinates of the center of bounding box No.1, along with its height and width. *C*_*i*_ is the label for the *i*^*th*^ class. For instance, in segmentation tasks, a dedicated head for segmentation is required, resulting in predictions being presented in a new format outlined in Formula (3). Instead of solely providing *x*, *y*, *w* and *h* for the bounding box, segmentation includes a mask comprising pairs of *x*, *y* coordinates that describe the boundaries of the target.

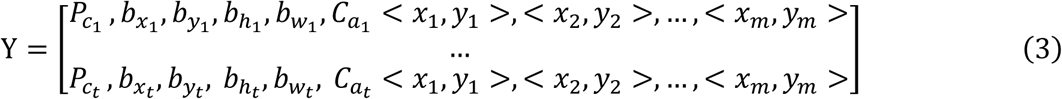

**Fig. S14** illustrates an example of *S* = 3, denoting a grid with 3 × 3 squares. The left side shows labeled objects in different shapes, while the corresponding feature vectors are shown on the right. The first element in each vector represents the probability of object existence. If it is 0, we disregard the remaining parts marked with “?”, as indicated by the gray area in the image. The next four values define the parameters for bounding boxes, followed by class labels representing AMR detection results, where orange denotes positive and green denotes negative. A blue polygon outlines the segmented boundaries, with an additional set of coordinate pairs within the feature vector defining the mask.

### 4.8. YOLO model fine-tuning

Object detection and segmentation are crucial functions in the field of computer vision, with applications from facial recognition to autonomous driving systems. Several popular object detection algorithms exist, including the R-CNN series ^47^, YOLO series ^48^, MobileNet ^49^, and SSD series ^50^. The YOLO (You Only Look Once) algorithm can achieve rapid detection. While it demonstrates impressive accuracy on the ImageNet object detection benchmark, it is limited to recognize objects within its predefined categories. Fine-tuning pre-trained models with a customized dataset that includes new object classes can enhance their capabilities.

The fine-tuning pipeline included four major steps: (i) preparation of the annotated dataset, (ii) fine-tuning environment configuration, (iii) model training and optimization, and (iv) evaluation. During training, we adopted the YOLOv5 architecture with the detection head frozen to preserve pretrained feature representations while adapting the model to new classes. The training hyperparameters were set as follows: learning rate 0.005 with cosine decay schedule, batch size 8, weight decay 0.0005, and momentum 0.937. Training was conducted for 500 epochs using stochastic gradient descent (SGD) as the optimizer.

For tasks such as object detection and segmentation, a key aspect of optimization is minimizing a loss function that balances localization and classification. Localization loss measures the error in predicting the location of an object, while classification loss measures the error in predicting the category of the object. By optimizing these loss functions particularly in a deep learning context, we improved performance in both dimensions, namely localization and classification. We used a loss function that includes bounding box regression loss (Bbox loss), objectness loss, and classification loss. We focused on the detection task as an example. The loss function is expressed as Formulas (4) and (5):

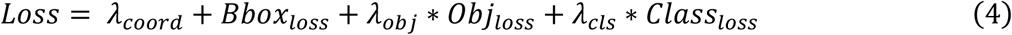

*loss*

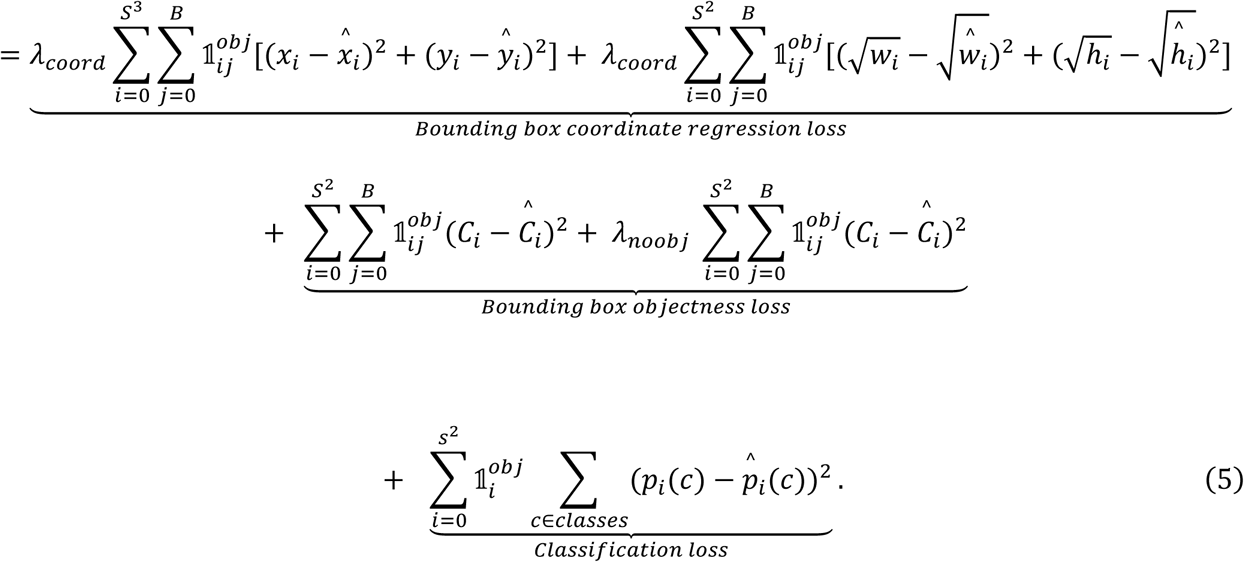

*S* represents the dividing parameter of grid. B denotes the number of bounding boxes per grid. The centroid location of each bounding box is given by *x*_*i*_, *y*_*i*_, and its dimensions are defined by *w*_*i*_(width) and *h*_*i*_ (height). *C*_*i*_ refers to objectness or confidence score, indicating whether an object exists. *P*_*i*_(*c*) signifies the classification loss. We applied masks 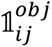 and 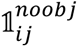 to each grid well. If a well in the microfluidic chip contains an object, 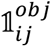 is set to 1; otherwise, it is 0. In contrast, 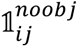 is set to 1 when no object is present and 0 when an object is in the well. The bounding box loss quantifies the difference between the predicted and actual bounding boxes. YOLO uses the Generalized Intersection over Union (GioU) loss ^51^, an improved version of the Intersection over Union (IoU) loss that also accounts for the area outside the union of the boxes. The formula for GioU is:

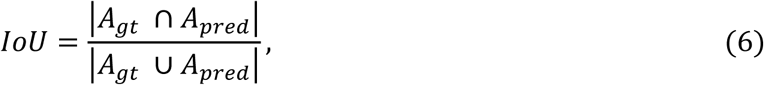

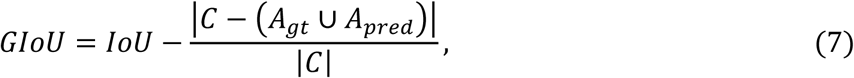

, where *A*_*gt*_ is the area of the ground truth box, *A*_*pred*_ is the area of the predicted box, and *C* is the area of the minimum bounding box containing the two boxes. The objectness loss measures the difference between the predicted and actual presence of an object within a bounding box. This is computed using Binary Cross-Entropy (BCE) loss. Its formula is:

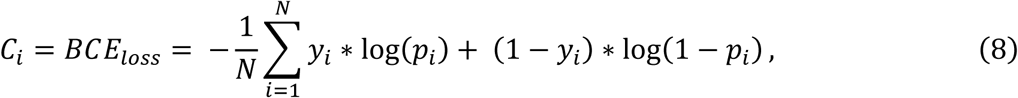

where *y*_*i*_ is the actual object presence (0 or 1) and *p*_*i*_ is the predicted object presence for sample *i*. It can be a decimal between 0 and 1. The classification loss represents the difference between the predicted and actual classes of an object within a bounding box. Cross-entropy is commonly used for this. Its formula is:

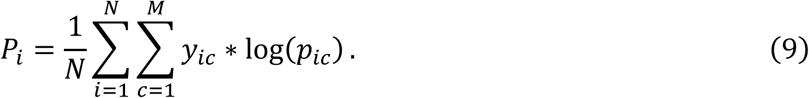

Similar to BCE loss, cross-entropy loss is calculated for each category individually. Here, *y*_*ic*_represents the ground truth value and *p*_*ic*_denotes the predicted probability that sample *i* belongs to category *c*.

The loss function plays a critical role in fine-tuning our model, guiding it to accurately identify AMR results from the microfluidic chips. By optimizing the loss function, the model improves its ability to recognize both the target category and the location of new targets, including well positions and AMR testing results. However, minimizing the loss function alone does not offer a comprehensive evaluation of the performance of the model. To assess the quality of detection by the model, we used additional metrics for evaluation alongside the loss function, including Precision [Formula (10)], Recalls [Formula (11)], and Mean Average Precision (mAP) [Formula (12)]. The mAP is the average of the Average Precision (AP) values calculated across all classes and recall levels. In object detection, AP represents the area under the Precision-Recall curve.

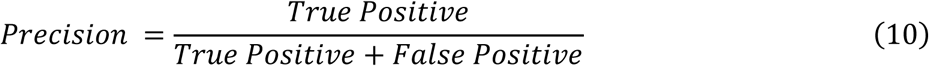

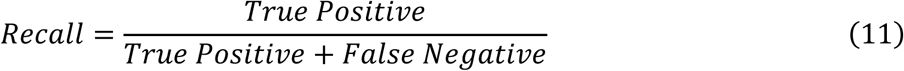

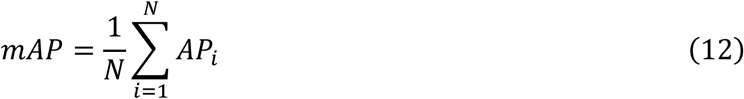

AP is a score ranging from 0 to 1, representing the area under the Precision-Recall Curve. Directly calculating this area can be impractical so that approximation methods (e.g., 11-point interpolation method) are often used. This method involves selecting 11 fixed recall thresholds, calculating maximum precision at each threshold, and averaging these values to determine AP as shown in Formula (13). *P*_*interp*_(*r*) represents the interpolated precision, which is the highest precision among all predicted values with a recall equal to or exceeding *r*.

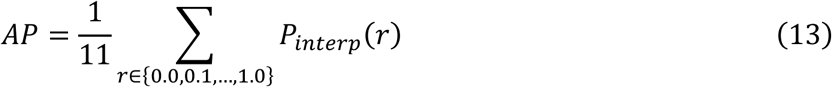

### 4.9. Model deployment strategies

Practical application of AIoT system is time sensitive in industrial settings with large-scale detection tasks. To address this, various YOLO model compression techniques can be employed to improve detection speed and maintain high accuracy. Common methods for model compression include quantization and pruning ^52^. Quantization reduces the numerical precision of the weights of a model, thereby reducing its size and inference time. Pruning involves removing less important weights or neurons to simplify the model without compromising performance. For quantization, a weight *w* is reduced to *Q*(*w*) using a function *Q*, with accuracy loss defined as |*w* − *Q*(*w*)|. Pruning typically uses a threshold *θ*. If the absolute value of a weight is below this threshold (|*w*| < *θ*), it is pruned to 0. These compression techniques allow AIoT systems to achieve fast detection, meeting industrial demands for speed and scalability. In our approach, we used a quantization strategy known as post-training quantization (PTQ). The original model was fine-tuned and formatted in Float 32 byte and then converted to Int8 format using the PTQ algorithm. This conversion reduced the size of the model by 80% and significantly improved inference speed by 1.5× with minimal effect on accuracy.

To deploy the optimized models onto a new platform, such as transitioning from an ×86-64 (AMD64) to an ARM64 platform, cross-compilation is necessary. This process involves adapting the runtime files and configurations of the model for compatibility with Orange Pi 5B. We used open-source software from Rockchip company to deploy the model on Rockchip chips, which power Orange Pi 5B’s SoC RK3588s. To leverage the built-in NPU of Orange Pi 5B that has a computing power of 6Tops@Int8, we first converted the PyTorch-trained “.pt” model to ONNX format using the Open Neural Network Exchange (ONNX) tool on an ×86 computer ^53^. Next, we used RKNN-Toolkit2 running on an ×86 platform to convert the ONNX model into RKNN format, which is compatible with Rockchip’s NPU. Finally, inference was performed through the provided Python API on the Orange Pi 5B.

### 4.10. Visualization interface of AMR surveillance on the cloud platform

The AMR surveillance platform was deployed on an Ubuntu operating system using a front-end/back-end separation architecture. The front end was developed with the Next.js framework, while the back end ran on Node.js v20.18.3. The back-end API service was implemented with the Express framework, and data were stored in MongoDB v8.0.5, accessed through the MongoDB Node.js driver (mongodb@6.16.0). RESTful APIs were employed to enable efficient communication between the front and back ends. The visualization interface integrated charting libraries such as ECharts and Mapbox to provide dynamic displays of data statistics, trend analyses, geographic distributions, and alert notifications. Fully deployed on cloud servers, the system offers scalability, high availability, and reliable technical support to enhance AMR surveillance and data analysis.

## Code availability

The code used in this study is available on GitHub (https://github.com/scaactk/AIoT_AMR).

## Acknowledgement

This work was supported by Canada Research Chairs Program (Grant # CRC-2024-00011), the Natural Sciences and Engineering Research Council of Canada in the form of a Discovery Grant (RGPIN-2025-03996), Agriculture and Agri-Food Canada (AAFC) AgriRisk Initiatives, Mitacs Accelerate, and Fonds de recherche du Québec-Nature et technologies (FRQNT) Strategic Clusters. We thank Dr. Douglas Call and Lindsay Parrish from Washington State University for providing some *Salmonella* isolates, and Yao Xuan Seah and Jia Jing Heng from the National University of Singapore for their assistance with data annotation.

## Supplementary Information

**Fig. S1.**
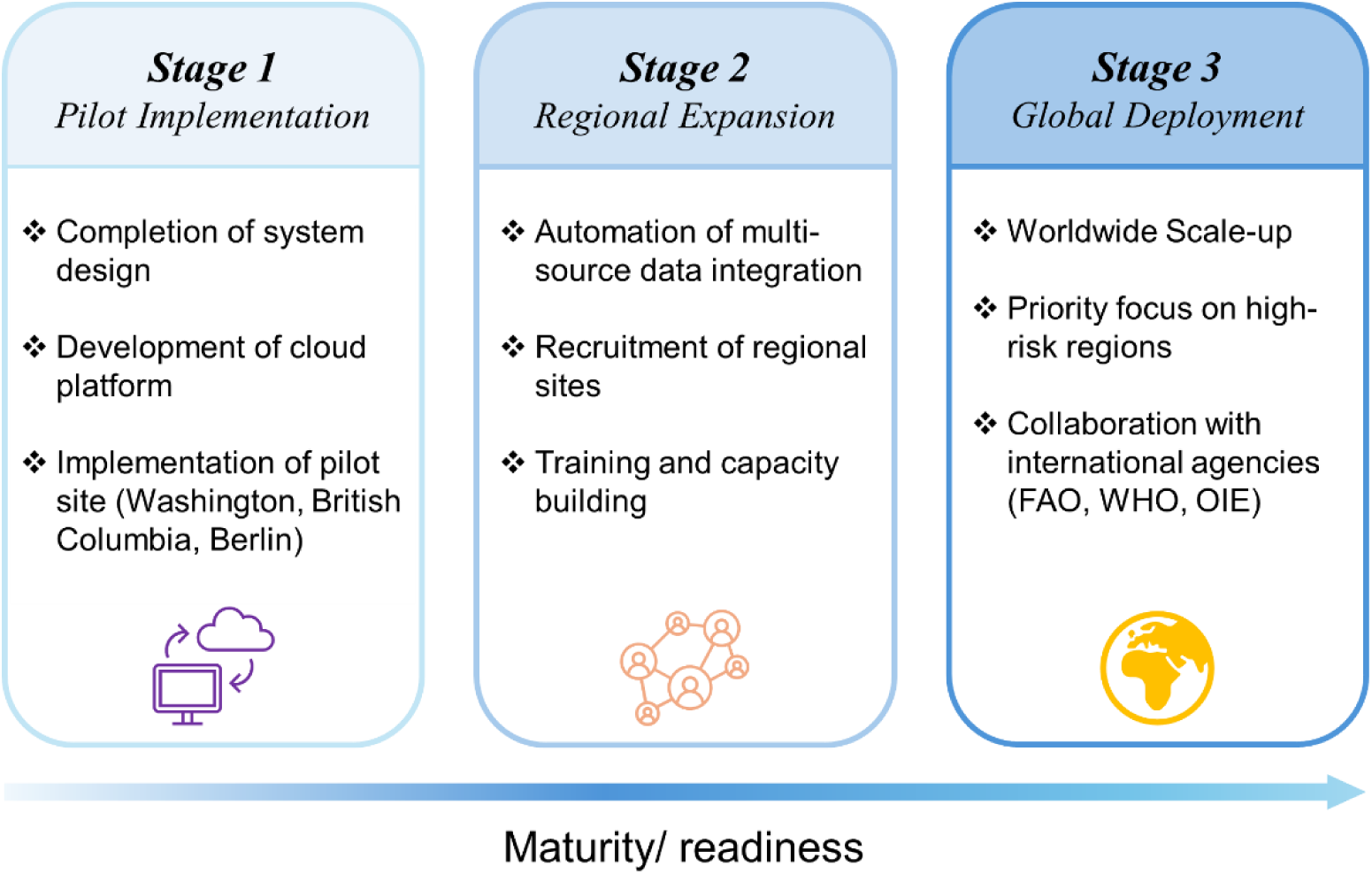
Development roadmap of the global AMR monitoring system for food supply chain. The roadmap outlines three stages of system evolution. ***Stage 1 (Pilot Implementation)*** encompasses system design, development of cloud platform, and initial pilot deployment at sites in Washington State (USA), British Columbia (Canada), and Berlin (Germany). ***Stage 2 (Regional Expansion)*** focuses on automating multi-source data integration, expanding regional surveillance networks, and strengthening capacity through training. ***Stage 3 (Global Deployment)*** represents worldwide scale-up, prioritizing high-risk regions and fostering collaboration with international organizations such as Food and Agriculture Organization (FAO), World Health Organization (WHO), and World Organization for Animal Health (WOAH). Collectively, these stages delineate the progression from pilot implementation to a fully operational global AMR surveillance network.

**Fig. S2.**
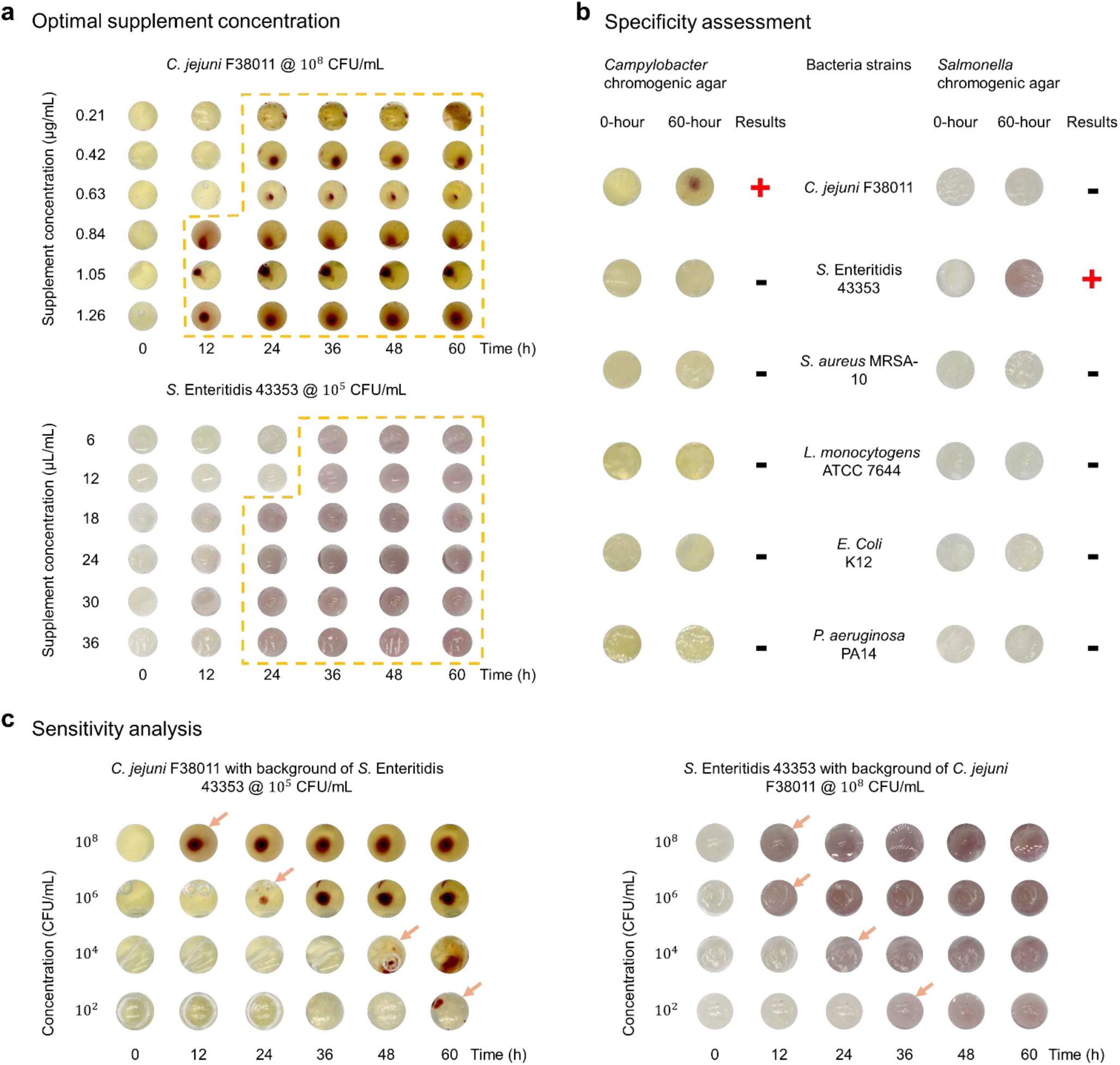
Testing and optimization of chromogenic agar for on-chip detection and differentiation of *Campylobacter* and *Salmonella*. a) Optimization of supplement concentrations in *Campylobacter* chromogenic agar using *C. jejuni* F38011, and in *Salmonella* chromogenic agar using *S.* Enteritidis 43353. b) Specificity testing of chromogenic agar with six bacterial strains (*C. jejuni* F38011, *S.* Enteritidis 43353, *S. aureus* MRSA-10, *L. monocytogenes* ATCC 7644, *E. coli* K12, and *P. aeruginosa* PA14) after incubation at 42°C for 60 h. c) Sensitivity testing with serial dilutions (10²–10⁸ CFU/mL) of *C. jejuni* F38011 mixed with *S.* Enteritidis 43353 at 10⁵ CFU/mL (left), and serial dilutions (10²–10⁸ CFU/mL) of *S.* Enteritidis 43353 mixed with *C. jejuni* F38011 at 10⁸ CFU/mL (right). All samples were incubated at 42°C for 60 h. Detection limits were defined as the lowest concentration at which a visible color change was observed (indicated by arrows).

**Fig. S3.**
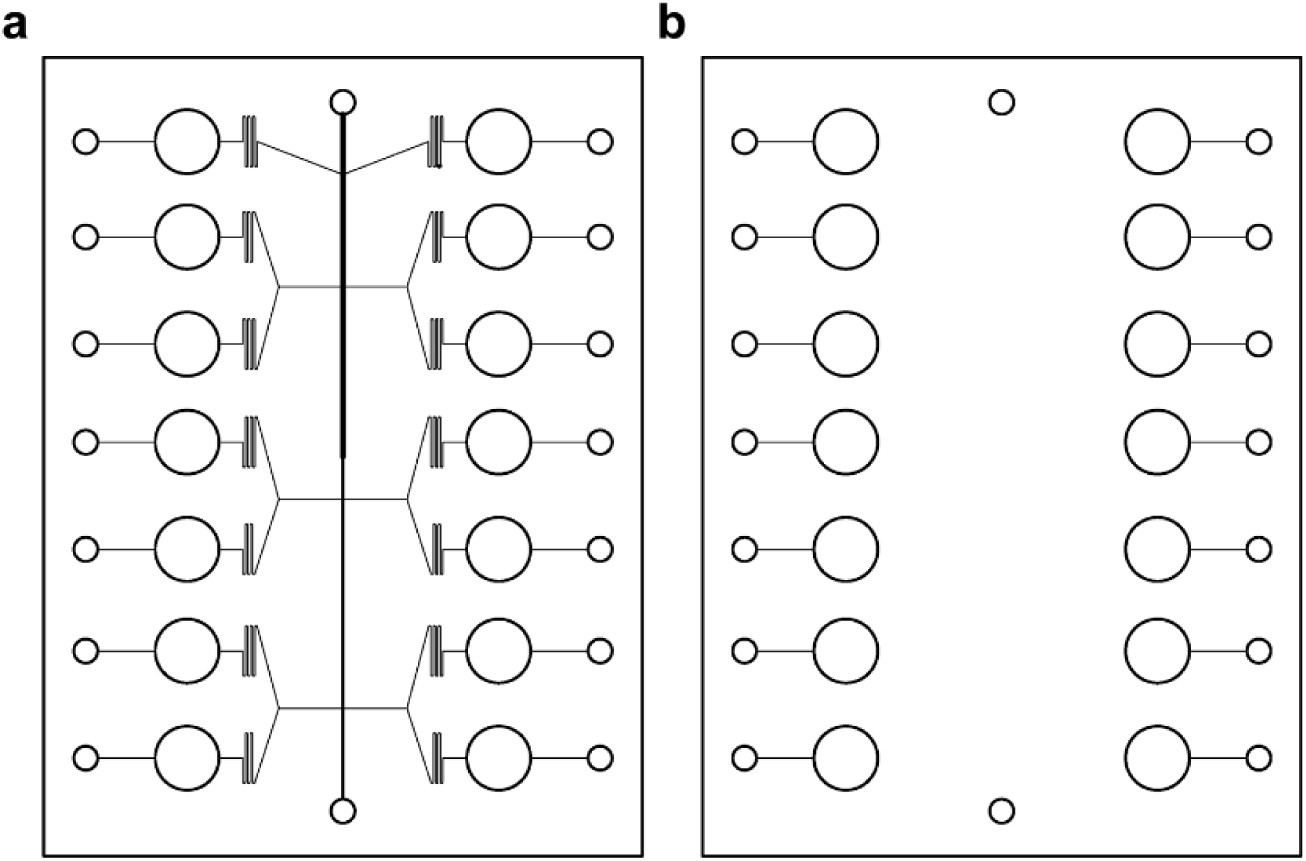
Layer configurations of the microfluidic chip. a) Schematic illustration of the middle layer depicting the microchannel network and reaction chamber arrangement for fluid distribution and sample processing. b) Layout of the bottom layer showing well positions and alignment holes to enable accurate assembly with the upper layers.

**Fig. S4.**
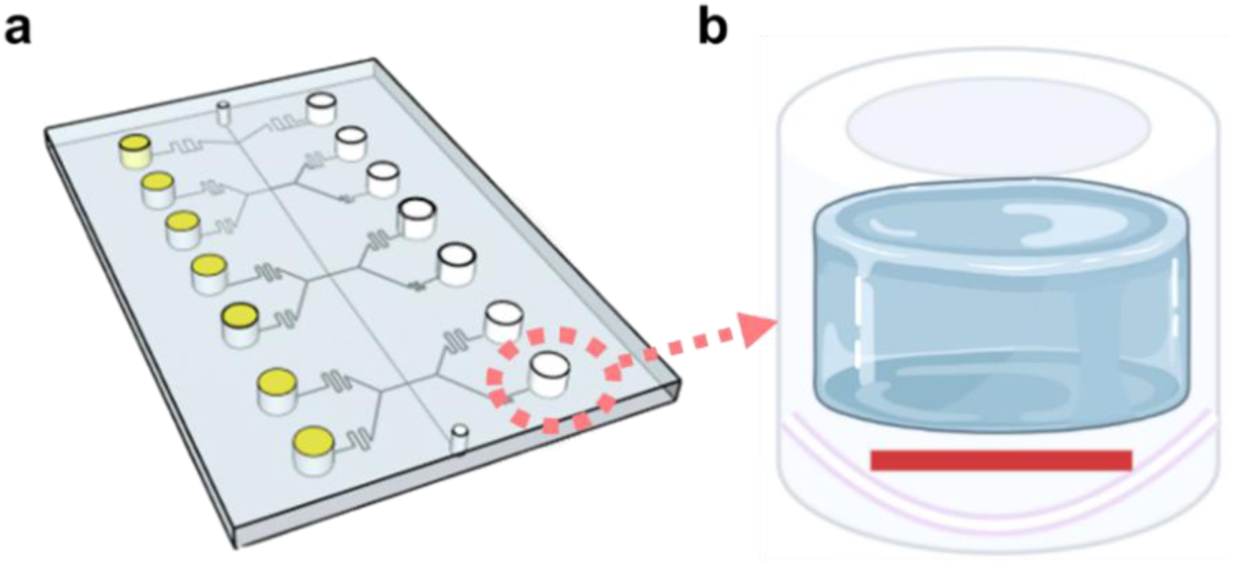
Diagram of the assembled microfluidic chip and chamber structure. a) Schematic illustration of the fully assembled microfluidic chip. Yellow chambers are filled with *Campylobacter* chromogenic agar, while white chambers contain *Salmonella* chromogenic agar. b) Cross-sectional view of a single reaction chamber. The purple-white line represents the PVDF membrane, the red line denotes the antimicrobial paper disk, and the blue, gel-like region indicates the agar.

**Fig. S5.**
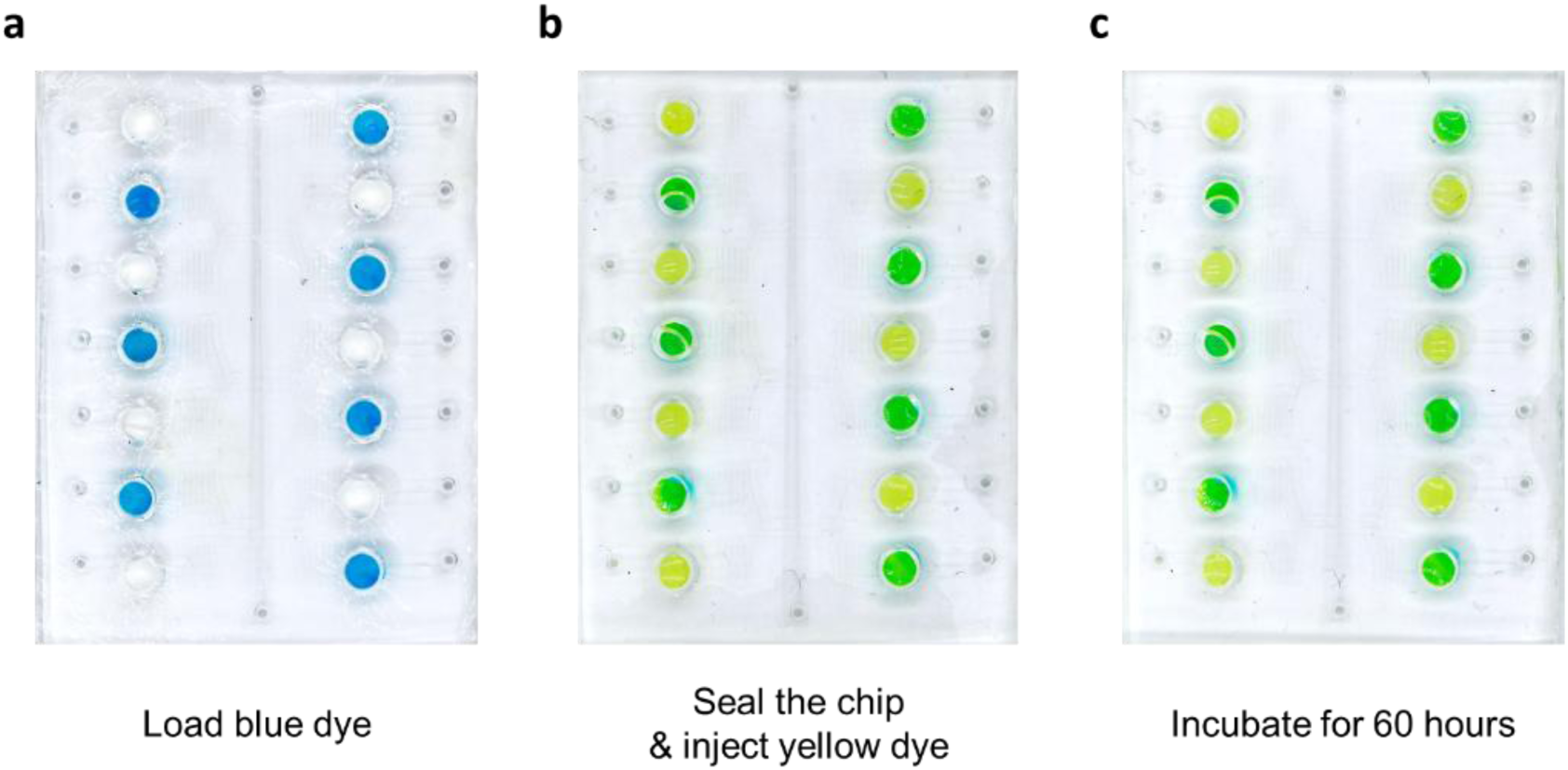
Evaluation of cross-contamination among incubation chambers. a) Paper disks containing either blue dye or left blank were placed in the incubation chambers of the microfluidic chip. b) Yellow dye solution was injected until all chambers were filled. Chambers containing blue dye turned green due to mixing, while the others remained yellow. a) To assess diffusion, the chip was incubated in a CO₂ incubator for 60 hours to simulate bacterial growth conditions. Chambers initially containing only yellow dye showed no color change, indicating no detectable diffusion from adjacent green chambers. These findings confirm that the chip design effectively prevents cross-contamination.

**Fig. S6.**
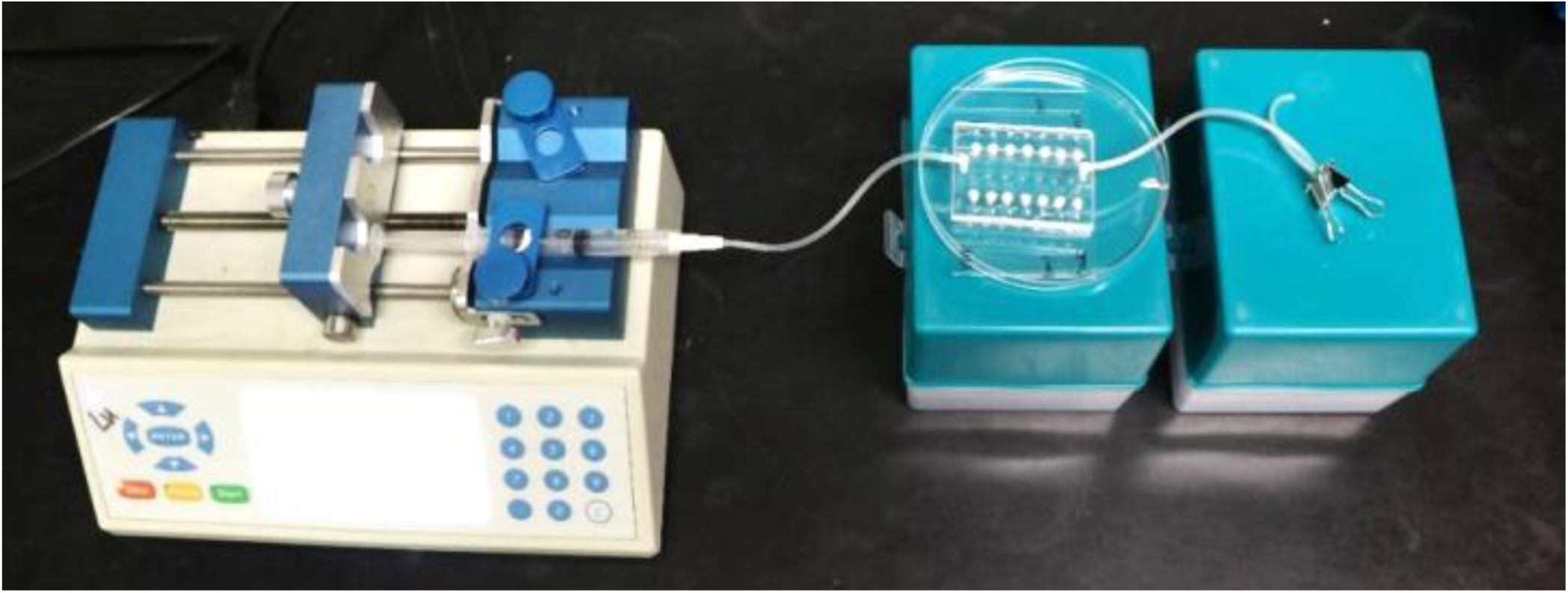
Setup for sample injection into the microfluidic chip. A syringe pump equipped with sterile accessories (syringe, tubing, and adapters) delivers the sample fluid into the microfluidic chip. Waste is collected in a Petri dish, while a binder clip placed on the outlet tubing serves as a stopper during sample loading.

**Fig. S7.**
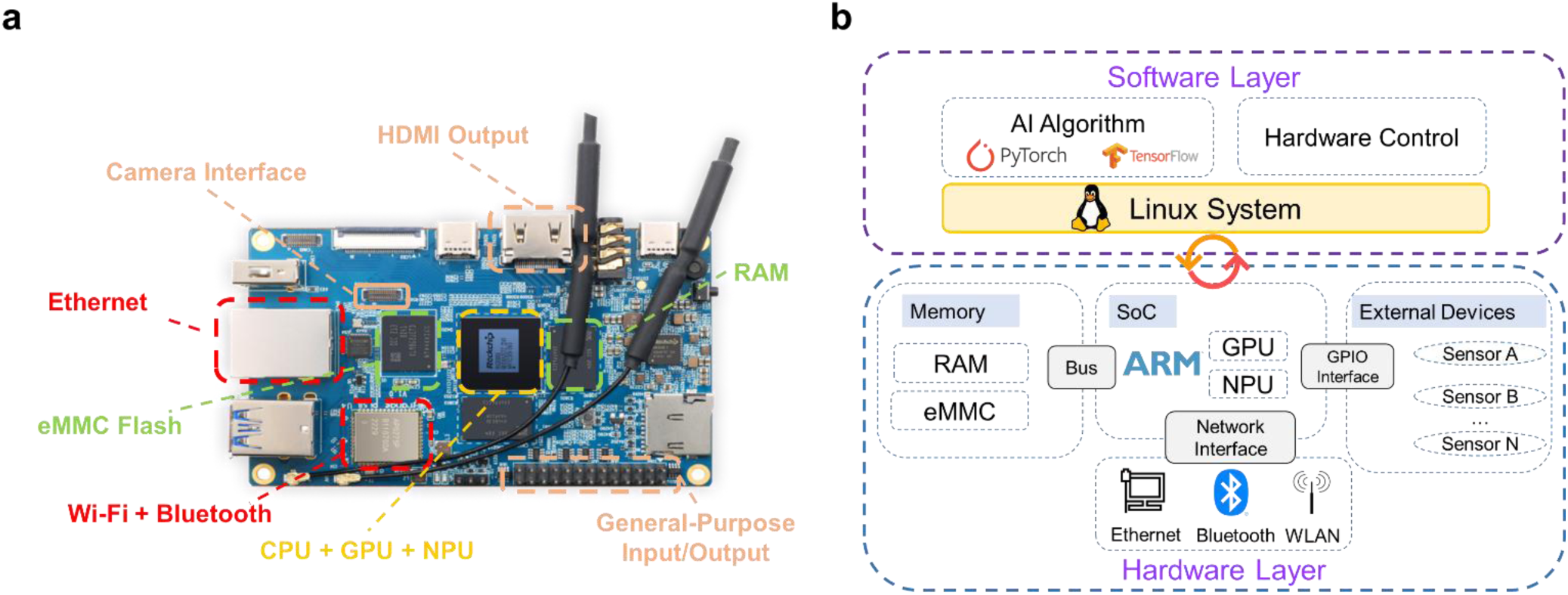
Functional structure of the Orange Pi system. a) Hardware layout of the Orange Pi 5B computer. The system-on-chip (SoC, including CPU, GPU, and NPU) is highlighted in yellow. Input/output components such as HDMI output, camera interface, and general-purpose input/output (GPIO) are marked in orange. Storage elements, including eMMC flash, are shown in green, while network-related components (Ethernet, Wi-Fi, and Bluetooth) are indicated in red. b) Integration of hardware and software within the Orange Pi system architecture. The hardware layer illustrates the interconnections among memory, SoC, bus, network interfaces, and external devices (e.g., sensors). The software layer depicts the hierarchical structure of the monitoring system, including the Linux operating system, AI frameworks, and custom hardware control strategies applied in this study.

**Fig. S8.**
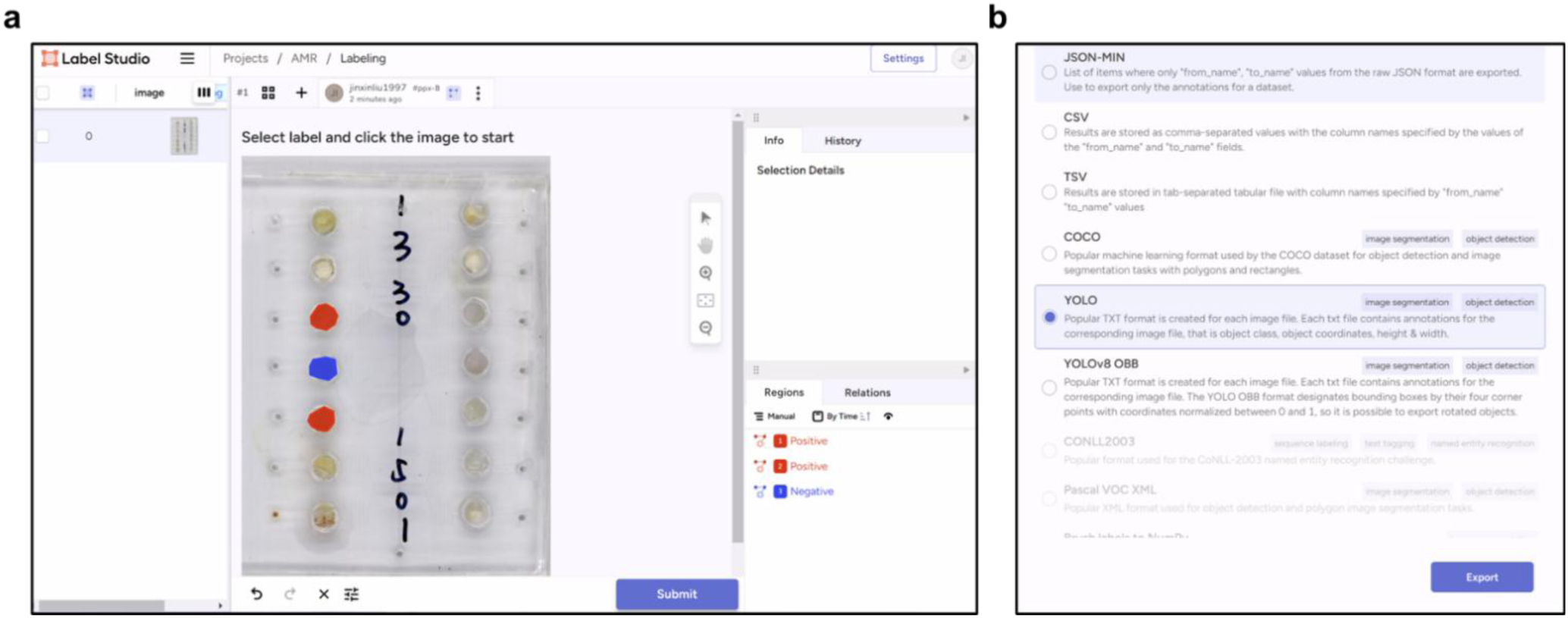
Label Studio image annotation and export interface. a) Image annotation interface in Label Studio, showing manual labeling of microfluidic chip wells as positive or negative classes. Annotators select labels and mark regions of interest (ROIs) directly on the image. b) Result export interface, which supports multiple annotation formats including YOLO, COCO, CSV, and others. The YOLO format is highlighted here for downstream machine learning applications.

**Fig. S9.**
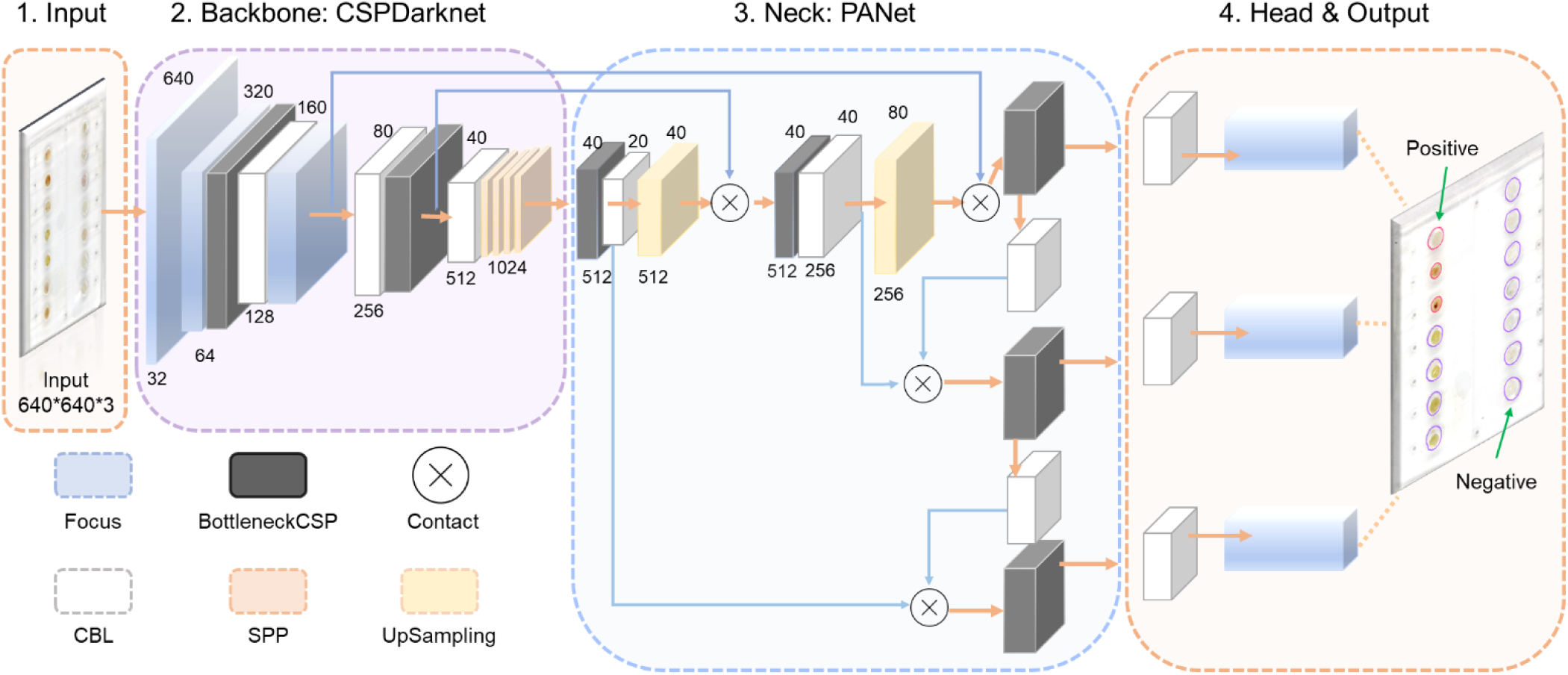
Structure of the YOLOv5 model. The YOLOv5 architecture consists of four main components: input, backbone, neck, and head. The leftmost section represents a 640 × 640 compressed image used as the network input. The backbone performs initial feature extraction, while the neck facilitates feature fusion to improve detection accuracy. The head generates tensors for specific tasks such as object detection and segmentation. After decoding, the output tensors correspond to markers at defined locations in the input image, distinguishing positive from negative signals.

### Some functional structures in YOLO v5

YOLO v5 incorporates several specialized functional structures that are crucial to its architecture, including **CBL** (Convolutional Block Lightweight), the **focus layer**, **SPP** (Spatial Pyramid Pooling), and **bottleneck CSP** (Cross Stage Partial). Below is an introduction to each component.

**CBL:** The CBL module is a lightweight block designed to replace traditional convolution layers. It typically consists of a convolution layer, a batch normal layer and an activation function sigmoid linear unit (SiLU), as shown in **Fig. S8**. Compared to traditional convolution layers, CBL reduces computational complexity and parameters required.

**Fig. S10.**
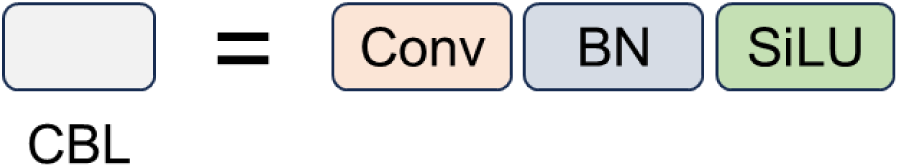
**Structure of CBL.**

**Focus layer**: The focus layer (**Fig. S9**) is a specialized convolutional structure that slices the input into four parts, each processed by a separate convolution operation. The results are then merged and passed through a CBL module. This slicing technique allows the focus structure to effectively capture both global and local information from the feature map.

**Fig. S11.**
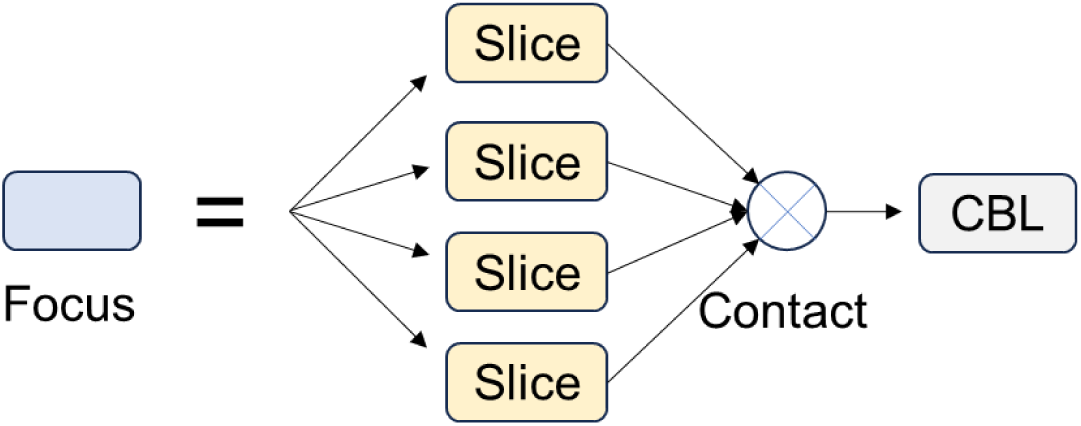
Structure of the focus layer.

**SPP**: The SPP module is a spatial pyramid pooling structure that enhances the robustness of the network to input scale variations and improves its receptive field (**Fig. S10**). The primary function of SPP is to maintain spatial information at multiple scales, enabling the network to recognize objects of different sizes within an image. SPP achieves this by applying parallel pooling operations with different grid configurations, typically involving max-pooling layers with varying kernel sizes and strides. In our neural network, the SPP module begins with a CBL block, followed by three Max Pooling blocks with different settings. The outputs from these Max Pooling blocks are concatenated with the output from the initial CBL block and then passed through another CBL block.

**Fig. S12.**
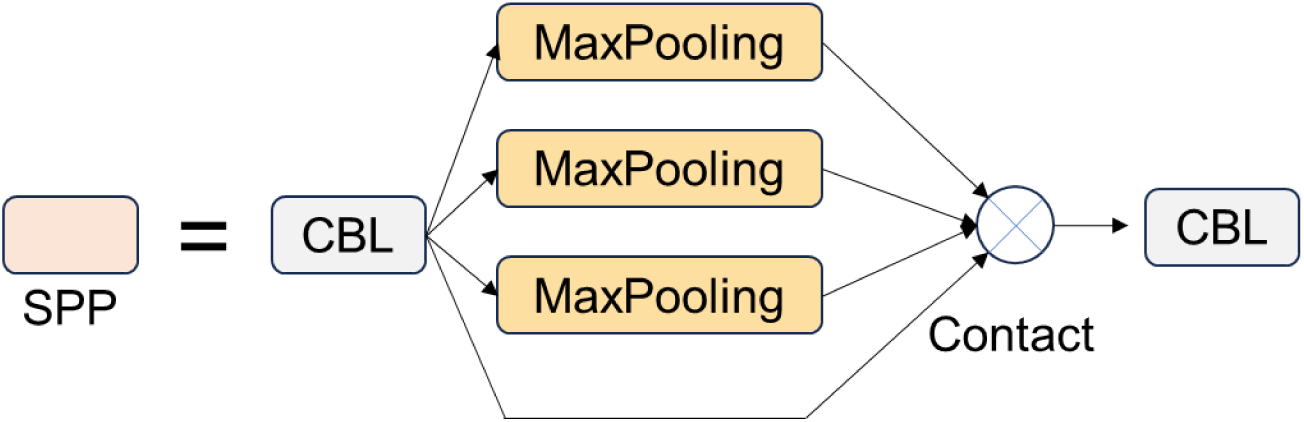
Structure of SPP block.

**Bottleneck CSP:** The Bottleneck CSP is a crucial component of YOLO v5, significantly enhancing the efficiency and performance of the model. Its detailed structure is shown in **Fig. S11**. Bottleneck CSP consists of two parts: the bottleneck and CSP. The bottleneck refers to a neural network design that reduces the dimensionality of input features through a sequence of layers. The key innovation in Bottleneck CSP is the use of partial connections across different stages of the network, allowing information to flow more freely between various parts, thereby improving feature propagation and representation learning.

**Fig. S13.**
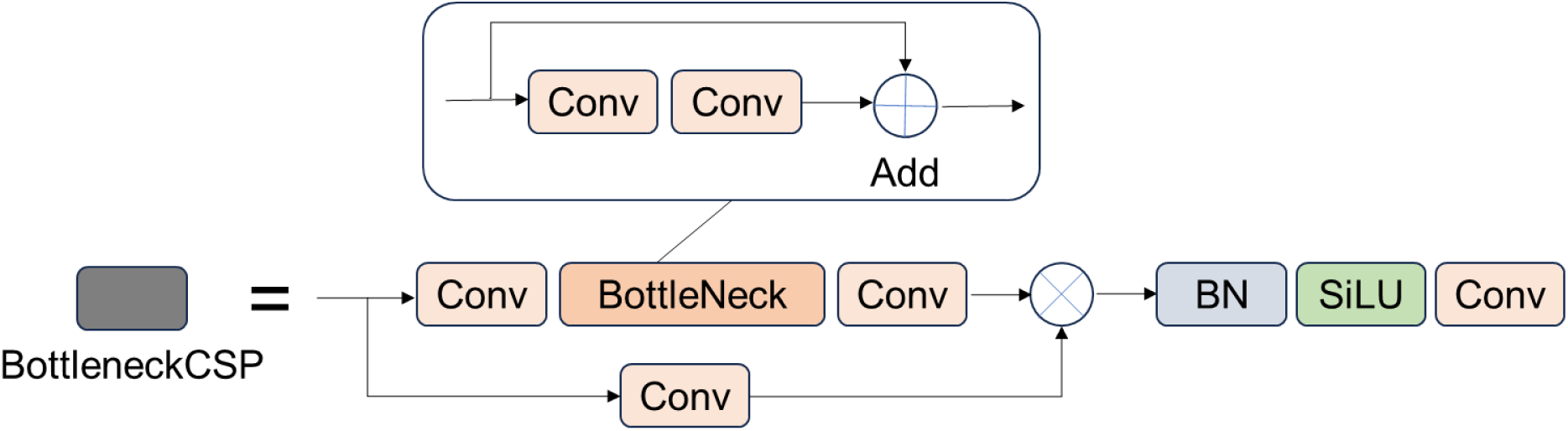
Structure of bottleneck CSP.

Incorporating these functional structures into YOLO v5 enhances its object detection capabilities, providing greater accuracy and efficiency. Proper design and integration of these components can significantly boost the performance of the model across various application scenarios.

**Fig. S14.**
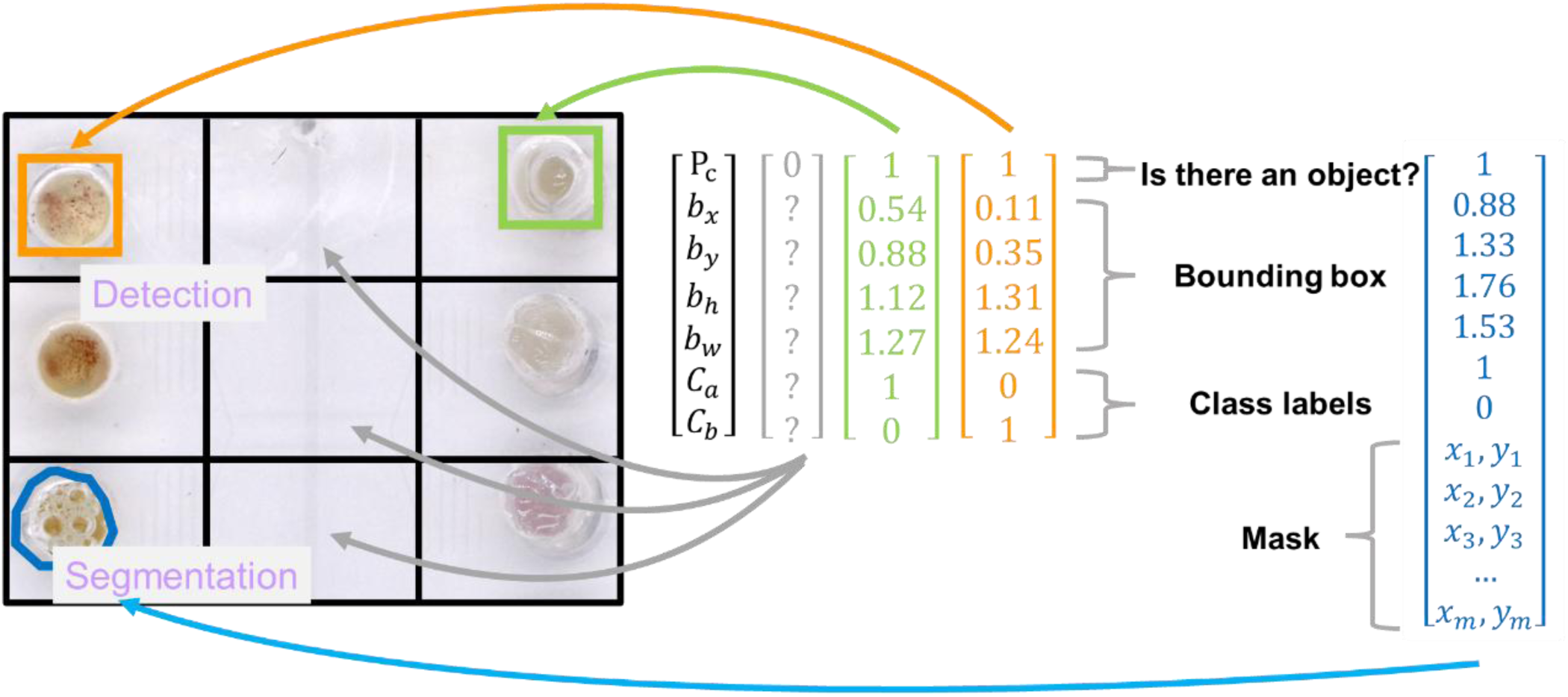
Illustration of YOLO detection and segmentation with corresponding feature vectors. Detection and segmentation outputs are shown on a 3 × 3 grid of microfluidic wells. Orange rectangles denote bounding boxes for positive targets, while green rectangles indicate negative targets. Blue polygons represent the segmented masks of detected targets. Each grid cell generates feature vectors that encode detection information (objectness and class scores) and segmentation information (mask coordinates).

**Fig. S15.**
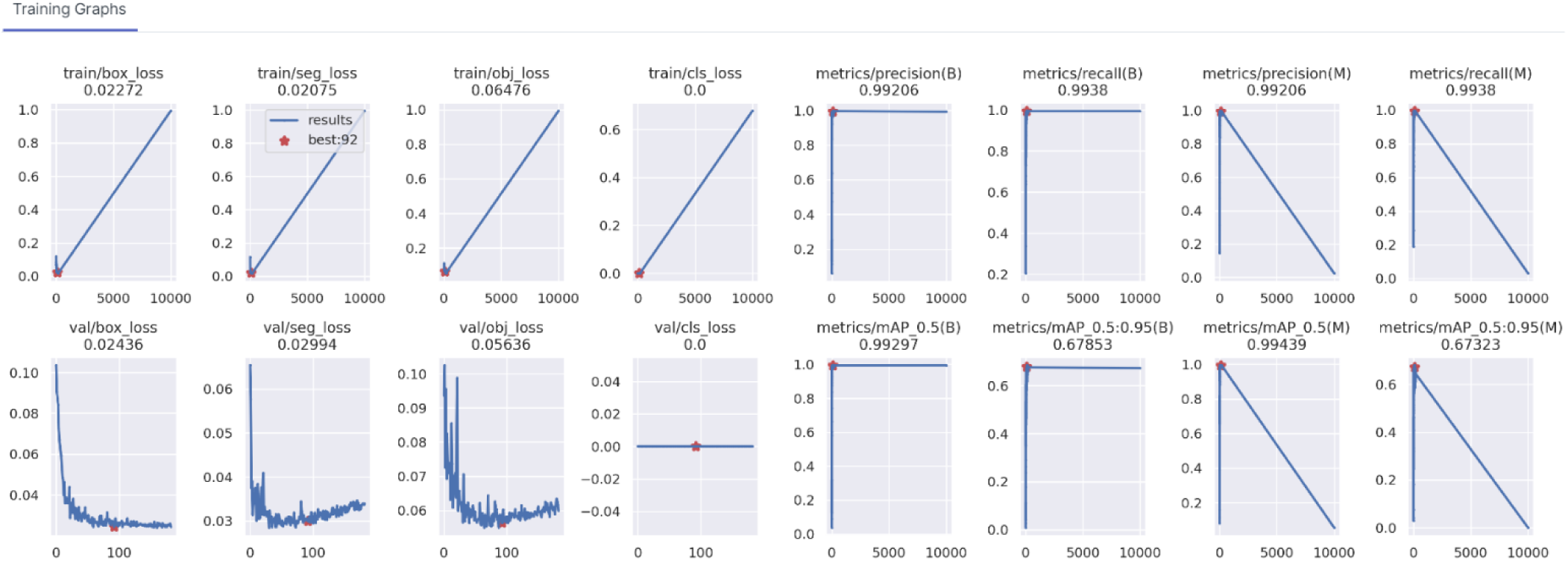
Example of model training and testing logs (detection). The training process tracks four loss functions: box_loss, seg_loss, obj_loss, and cls_loss. Evaluation metrics include precision, mean Average Precision (mAP), and recall. Indicators marked with (B) denote the best performance achieved during training, while (M) indicates the macro average across all classes. Specifically, mAP_0.5 refers to the mAP at an IoU threshold of 0.5, whereas mAP_0.5:0.95 represents the average mAP across IoU thresholds from 0.5 to 0.95.

**Table S1.** Minimum inhibitory concentration (MIC) breakpoints from CLSI for AMR test.

| *Campylobacter* | Susceptible breakpoint (µg/mL) | Resistant breakpoint (µg/mL) |
| --- | --- | --- |
| ciprofloxacin | ≤1 | ≥4 |
| erythromycin | ≤8 | ≥32 |
| tetracycline | ≤4 | ≥16 |
| *Salmonella* | Susceptible breakpoint (µg/mL) | Resistant breakpoint (µg/mL) |
| ampicillin | ≤8 | ≥32 |
| ciprofloxacin | ≤0.06 | ≥1 |
| *tetracycline* | ≤4 | ≥16 |

**Table S2.**
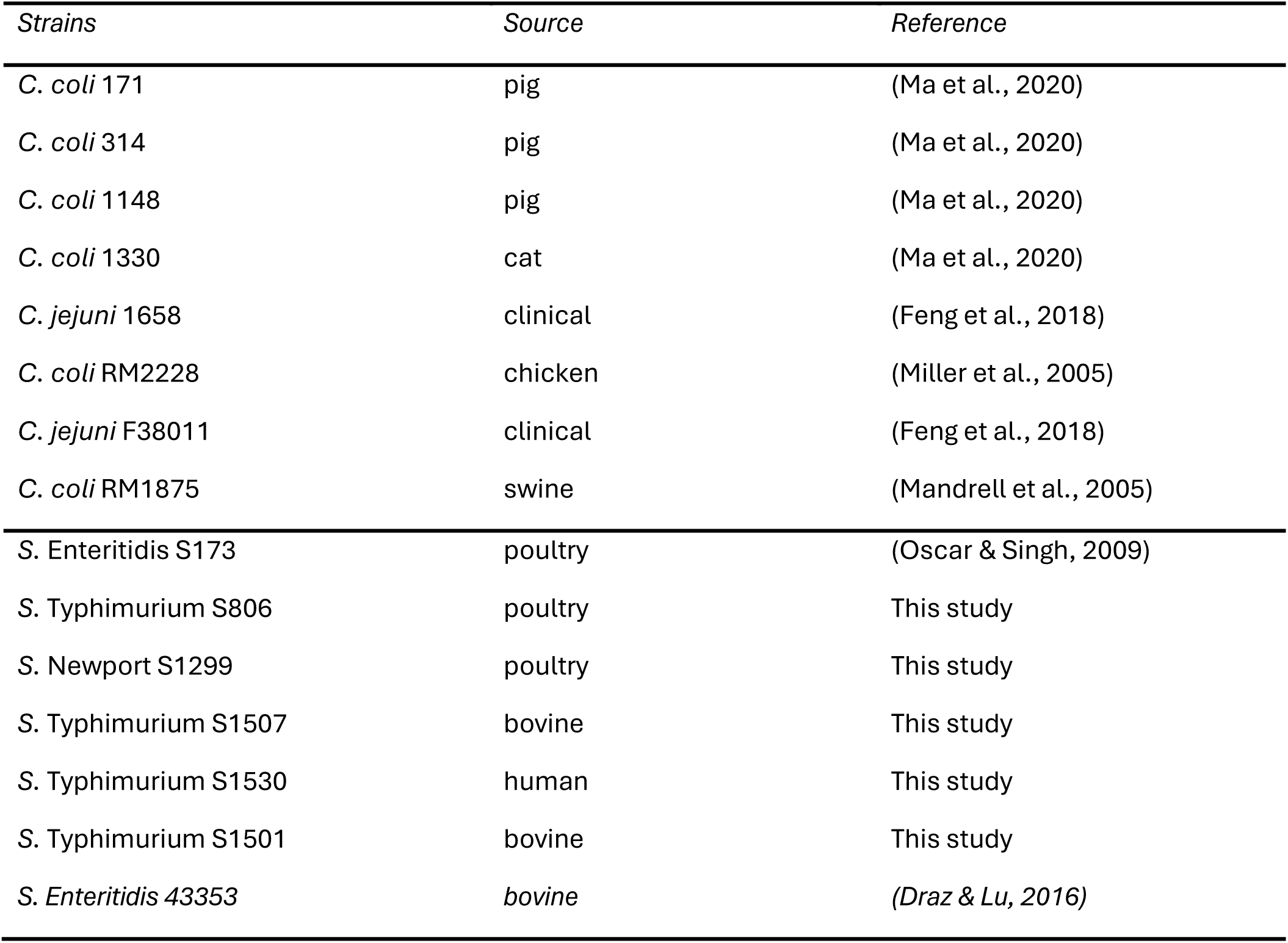
*Campylobacter* and *Salmonella* strains used for AST.

| *Strains* | *Source* | *Reference* |
| --- | --- | --- |
| *C. coli* 171 | pig | (Ma et al., 2020) |
| *C. coli* 314 | pig | (Ma et al., 2020) |
| *C. coli* 1148 | pig | (Ma et al., 2020) |
| *C. coli* 1330 | cat | (Ma et al., 2020) |
| *C. jejuni* 1658 | clinical | (Feng et al., 2018) |
| *C. coli* RM2228 | chicken | (Miller et al., 2005) |
| *C. jejuni* F38011 | clinical | (Feng et al., 2018) |
| *C. coli* RM1875 | swine | (Mandrell et al., 2005) |
| *S. Enteritidis* S173 | poultry | (Oscar & Singh, 2009) |
| *S. Typhimurium* S806 | poultry | This study |
| *S. Newport* S1299 | poultry | This study |
| *S. Typhimurium* S1507 | bovine | This study |
| *S. Typhimurium* S1530 | human | This study |
| *S. Typhimurium* S1501 | bovine | This study |
| *S. Enteritidis* 43353 | bovine | (Draz & Lu, 2016) |

**Table S3.**
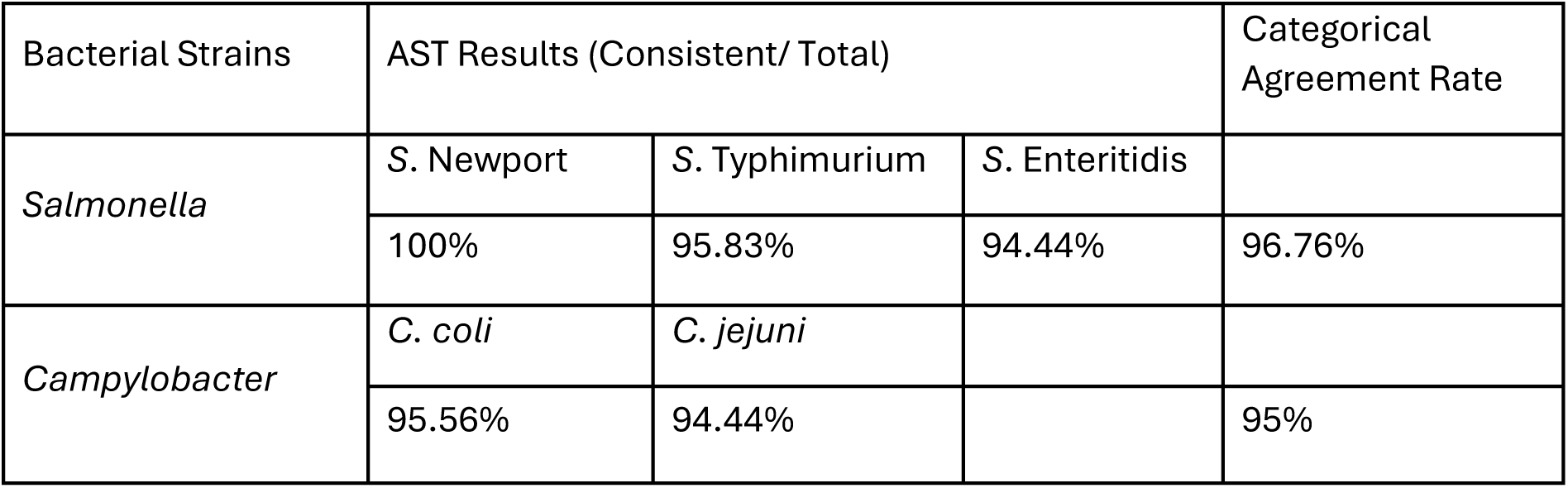
Agreement rates between the AIoT system and the standard AST method based on broth microdilution using locally isolated bacterial strains.

**Table S4.a.** Information on multiregional *Salmonella* isolates, MIC test results, and comparison with the AIoT-based on-chip testing system.

| Isolates ID & Group | Serovar | Ampicillin | Ciprofloxacin | Tetracycline | AloT Results Consistency |
| --- | --- | --- | --- | --- | --- |
| 1-B | Typhimurium | R | S | I | ☑ |
| 2-B | Typhimurium | R | S | R | ☑ |
| 3-B | Typhimurium | R | S | R | ☑ |
| 4-B | Saintpaul | R | S | R | ☑ |
| 5-B | Typhimurium | S | I | R | ☑ |
| 6-C2 | Kentucky | S | S | R | ☑ |
| 7-C2 | Kentucky | S | S | R | ☑ |
| 9-C2 | Kentucky | S | S | S | ☑ |
| 10-D1 | Enteritidis | S | S | S | ? |
| 11-D1 | Enteritidis | S | I | S | ☑ |
| Total agreement rate |  |  |  |  | 96.7% |

\* *Salmonella* isolates' ID are custom defined by its number and sample group.
\* "R" indicates Resistance, "S" indicates susceptible, and "I" indicates intermediate.
\* ☑ indicates complete agreement between AloT results and standard MIC based on broth microdilution.
\* "?" indicates that some results are inconsistent.

**Table S4.b.**
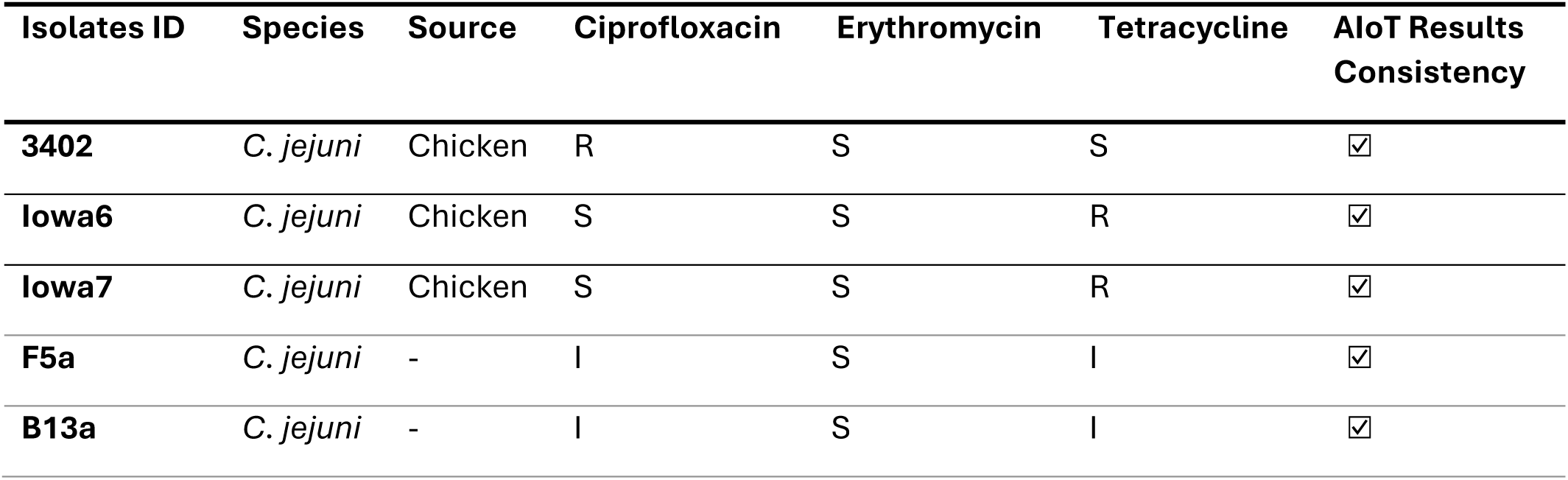
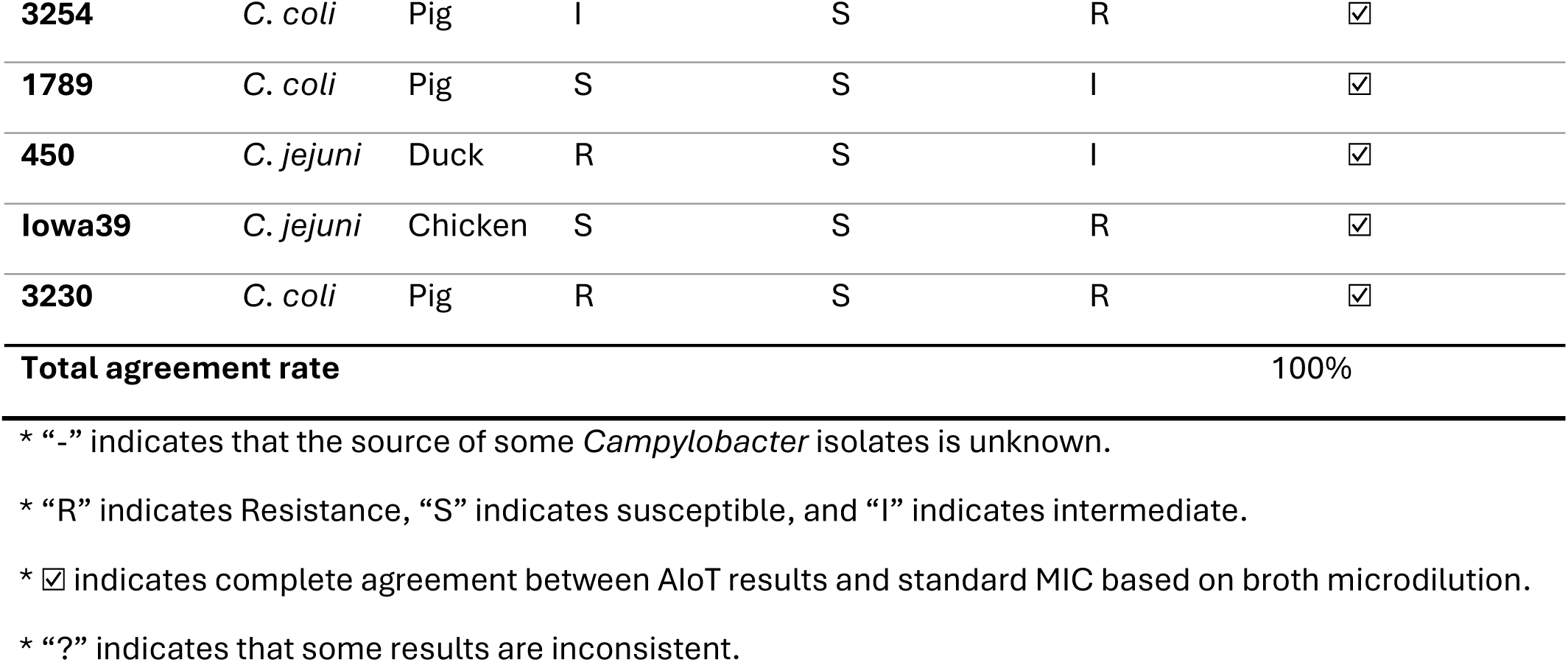
Information on multiregional *Campylobacter* isolates, MIC test results, and comparison with the AIoT-based on-chip testing system.

**Table S5.** Comparison of cost between AIoT-enhanced solution and the conventional AST method.

| Category | AIoT-enhanced solution |  | Conventional AST method |  |
| --- | --- | --- | --- | --- |
| Fixed cost | Homemade incubator box | \$15 | Commercial Solution | >\$10,000 |
| | AI system | \$100 | | |
| | Sensors | \$50 | | |
| Variable cost | Microfluidic Chip | \$ 0.06 per chamber | 96 well plate | \$0.15 per well |
|  | Growth media & antibiotics | Cheap (20 µL/test) | Growth media & antibiotics | Relatively pricy (200 µL/test) |

Prices are shown in USD.

## References

1. Salam, M. A. et al. Antimicrobial Resistance: A Growing Serious Threat for Global Public Health. Healthcare 11, 1946 (2023).

2. Aminov, R. I. A Brief History of the Antibiotic Era: Lessons Learned and Challenges for the Future. Frontiers in Microbiology 1, (2010).

3. Cook, M. A. & Wright, G. D. The past, present, and future of antibiotics. Science Translational Medicine 14, eabo7793 (2022).

4. O’Neill, J. Antimicrobial resistance: tackling a crisis for the health and wealth of nations. Rev. Antimicrob. Resist. (2014).

5. Sagar, P., Aseem, A., Banjara, S. K. & Veleri, S. The role of food chain in antimicrobial resistance spread and One Health approach to reduce risks. International Journal of Food Microbiology 391–393, 110148 (2023).

6. Thakur, S. & Gray, G. C. The Mandate for a Global “One Health” Approach to Antimicrobial Resistance Surveillance. Am J Trop Med Hyg 100, 227–228 (2019).

7. Sayad, A. et al. A microdevice for rapid, monoplex and colorimetric detection of foodborne pathogens using a centrifugal microfluidic platform. Biosensors and Bioelectronics 100, 96–104 (2018).

8. Jiang, Y., Wang, H., Li, S. & Wen, W. Applications of Micro/Nanoparticles in Microfluidic Sensors: A Review. Sensors 14, 6952–6964 (2014).

9. Ma, L., Petersen, M. & Lu, X. Identification and Antimicrobial Susceptibility Testing of *Campylobacter* Using a Microfluidic Lab-on-a-Chip Device. Applied and Environmental Microbiology 86, e00096–20 (2020).

10. Yang, J. et al. A self-powered microfluidic chip integrated with fluorescent microscopic counting for biomarkers assay. Sensors and Actuators B: Chemical 291, 192–199 (2019).

11. Coelho, J. R. et al. The Use of Machine Learning Methodologies to Analyse Antibiotic and Biocide Susceptibility in *Staphylococcus aureus*. PLOS ONE 8, e55582 (2013).

12. Pyayt, A., Khan, R., Brzozowski, R., Eswara, P. & Gubanov, M. Rapid Antibiotic Susceptibility Analysis Using Microscopy and Machine Learning. in 2020 IEEE International Conference on Big Data (Big Data) 5804–5806 (2020). doi:10.1109/BigData50022.2020.9378005.

13. Thrift, W. J. et al. Deep Learning Analysis of Vibrational Spectra of Bacterial Lysate for Rapid Antimicrobial Susceptibility Testing. ACS Nano 14, 15336–15348 (2020).

14. Madakam, S., Ramaswamy, R. & Tripathi, S. Internet of Things (IoT): A Literature Review. Journal of Computer and Communications 3, 164–173 (2015).

15. Chi, H. R., Wu, C. K., Huang, N.-F., Tsang, K.-F. & Radwan, A. A Survey of Network Automation for Industrial Internet-of-Things Toward Industry 5.0. IEEE Transactions on Industrial Informatics 19, 2065–2077 (2023).

16. Saranya, T., Deisy, C., Sridevi, S. & Anbananthen, K. S. M. A comparative study of deep learning and Internet of Things for precision agriculture. Engineering Applications of Artificial Intelligence 122, 106034 (2023).

17. Liang, Y. et al. UrbanFM: Inferring Fine-Grained Urban Flows. in Proceedings of the 25th ACM SIGKDD International Conference on Knowledge Discovery & Data Mining 3132–3142 (Association for Computing Machinery, New York, NY, USA, 2019). doi:10.1145/3292500.3330646.

18. Wang, S. et al. DelvMap: Completing Residential Roads in Maps Based on Couriers’ Trajectories and Satellite Imagery. IEEE Transactions on Geoscience and Remote Sensing 62, 1–14 (2024).

19. Dankan Gowda, V., et al. Dynamic Disaster Management with Real-Time IoT Data Analysis and Response. in 2024 International Conference on Automation and Computation (AUTOCOM) 142–147 (2024). doi:10.1109/AUTOCOM60220.2024.10486101.

20. Prapti, D. R. et al. Internet of Things (IoT)-based aquaculture: An overview of IoT application on water quality monitoring. Reviews in Aquaculture 14, 979–992 (2022).

21. Wang, K. et al. An RFID-based automated individual perching monitoring system for group-housed poultry. Transactions of the ASABE 62, 695–704 (2019).

22. Cocco, L. et al. A Blockchain-Based Traceability System in Agri-Food SME: Case Study of a Traditional Bakery. IEEE Access 9, 62899–62915 (2021).

23. Sourav, A. I., Lynn, N. D. & Suyoto. Smart Monitoring System Design for Perishable Food Supply Chain Management Based on IoT in Bangladesh. International Journal of Advanced Science and Technology 29, 1069–1079 (2020).

24. Ma, L., He, W., Petersen, M., Chou, K. C. & Lu, X. Next-Generation Antimicrobial Resistance Surveillance System Based on the Internet-of-Things and Microfluidic Technique. ACS Sens. 6, 3477–3484 (2021).

25. Shi, W., Cao, J., Zhang, Q., Li, Y. & Xu, L. Edge Computing: Vision and Challenges. IEEE Internet Things J. 3, 637–646 (2016).

26. Battersby, T., Walsh, D., Whyte, P. & Bolton, D. J. *Campylobacter* growth rates in four different matrices: broiler caecal material, live birds, Bolton broth, and brain heart infusion broth. Infection Ecology & Epidemiology 6, 31217 (2016).

27. Simner, P. J. et al. What’s New in Antibiograms? Updating CLSI M39 Guidance with Current Trends. J Clin Microbiol 60, e02210–21 (2022).

28. Shorten, C. & Khoshgoftaar, T. M. A survey on Image Data Augmentation for Deep Learning. J Big Data 6, 60 (2019).

29. Kirillov, A. et al. Segment anything. in Proceedings of the IEEE/CVF International Conference on Computer Vision 4015–4026 (2023).

30. Qin, D., Xia, Y. & Whitesides, G. M. Soft lithography for micro- and nanoscale patterning. Nat Protoc 5, 491–502 (2010).

31. Hunt, J. M., Abeyta, C. & Tran, T. BAM Chapter 7: *Campylobacter*. FDA (2024).

32. Andrews, W. H., et al. BAM Chapter 5: *Salmonella*. FDA (2024).

33. Kou, R. et al. Infrared Small Target Tracking Algorithm via Segmentation Network and Multistrategy Fusion. IEEE Transactions on Geoscience and Remote Sensing 61, 1–12 (2023).

34. Tan, T. & Cao, G. Efficient Execution of Deep Neural Networks on Mobile Devices with NPU. in Proceedings of the 20th International Conference on Information Processing in Sensor Networks (co-located with CPS-IoT Week 2021) 283–298 (ACM, Nashville TN USA, 2021). doi:10.1145/3412382.3458272.

35. Wootton, C. General Purpose Input/Output (GPIO). in Samsung ARTIK Reference 235–288 (Apress, Berkeley, CA, 2016). doi:10.1007/978-1-4842-2322-2_17.

36. Chen, J. & Huang, S. Analysis and Comparison of UART, SPI and I2C. in 2023 IEEE 2nd International Conference on Electrical Engineering, Big Data and Algorithms (EEBDA) 272–276 (2023). doi:10.1109/EEBDA56825.2023.10090677.

37. Nevliudov, I. et al. Mobile Robot Navigation System Based on Ultrasonic Sensors. in 2023 IEEE XXVIII International Seminar/Workshop on Direct and Inverse Problems of Electromagnetic and Acoustic Wave Theory (DIPED) vol. 1 247–251 (2023).

38. Xiong, W., Wang, Z., Zhang, B. & Li, S. Robust Voltage Regulation for DC–DC Converters via a Predictive GPIO-Based Control Approach. IEEE Transactions on Circuits and Systems II: Express Briefs 69, 4864–4868 (2022).

39. Paszke, A. et al. Pytorch: An imperative style, high-performance deep learning library. Advances in neural information processing systems 32, (2019).

40. Abadi, M. et al. TensorFlow: a system for Large-Scale machine learning. in 12th USENIX symposium on operating systems design and implementation (OSDI 16) 265–283 (2016).

41. Miell, I. & Sayers, A. Docker in Practice. (Simon and Schuster, 2019).

42. Yoshida, K., Miwa, S., Yamaki, H. & Honda, H. Analyzing the impact of CUDA versions on GPU applications. Parallel Computing 120, 103081 (2024).

43. Chetlur, S. et al. cuDNN: Efficient Primitives for Deep Learning. Preprint at http://arxiv.org/abs/1410.0759 (2014).

44. Redmon, J., Divvala, S., Girshick, R. & Farhadi, A. You Only Look Once: Unified, Real-Time Object Detection. in 779–788 (2016).

45. Liu, S., Qi, L., Qin, H., Shi, J. & Jia, J. Path aggregation network for instance segmentation. In Proceedings of the IEEE conference on computer vision and pattern recognition 8759–8768 (2018).

46. Deng, J. et al. Imagenet: A large-scale hierarchical image database. in 2009 IEEE conference on computer vision and pattern recognition 248–255 (Ieee, 2009).

47. He, K., Gkioxari, G., Dollár, P. & Girshick, R. Mask r-cnn. in Proceedings of the IEEE international conference on computer vision 2961–2969 (2017).

48. Jiang, P., Ergu, D., Liu, F., Cai, Y. & Ma, B. A Review of Yolo algorithm developments. Procedia computer science 199, 1066–1073 (2022).

49. Howard, A. G., et al. MobileNets: Efficient Convolutional Neural Networks for Mobile Vision Applications. Preprint at 10.48550/arXiv.1704.04861 (2017).

50. Liu, W. et al. SSD: Single Shot MultiBox Detector. in Computer Vision – ECCV 2016 (eds Leibe, B., Matas, J., Sebe, N. & Welling, M.) vol. 9905 21–37 (Springer International Publishing, Cham, 2016).

51. Rezatofighi, H. et al. Generalized intersection over union: A metric and a loss for bounding box regression. in Proceedings of the IEEE/CVF conference on computer vision and pattern recognition 658–666 (2019).

52. Choudhary, T., Mishra, V., Goswami, A. & Sarangapani, J. A comprehensive survey on model compression and acceleration. Artif Intell Rev 53, 5113–5155 (2020).

53. Jajal, P. et al. Interoperability in Deep Learning: A User Survey and Failure Analysis of ONNX Model Converters. in Proceedings of the 33rd ACM SIGSOFT International Symposium on Software Testing and Analysis 1466–1478 (ACM, Vienna Austria, 2024). doi:10.1145/3650212.3680374.

